# Identifying cytokine-release signatures of flow-driven endothelial remodelling in an intracranial aneurysm cell culture model

**DOI:** 10.64898/2026.06.18.733289

**Authors:** Chloe M. de Nys, Tiago Guerzet Sardenberg Lima, Haveena Anbananthan, Timothy Mitchell, Salma Mansi, Alyssa Binder, Zhiyong Li, James I. Novak, Petra Mela, Steven G. Wise, Danilo Carluccio, Craig D. Winter, Ashley R. Murphy, Rose Ann Franco, Mark C. Allenby

## Abstract

Intracranial aneurysm (IA) rupture is catastrophic, yet current models of rupture-risk inadequately capture underlying IA remodelling mechanisms. Endothelial-haemodynamic interactions are central to these processes, but *in vitro* flow platforms often lack vessel-relevant geometry or long-term perfusion. Here, temporal and spatial endothelial responses to haemodynamic stress were investigated across idealised and patient-specific vascular models.

Polydimethylsiloxane models were endothelialised with human aortic endothelial cells then perfused at up to 1.6 Pa wall shear stress for five days. IA models were exposed to steady or cardiovascular flow waveforms, with endothelial phenotype assessed by immunofluorescence and cytokine profiling.

Flow initiation induced a transient inflammatory response, with elevated MCP-1 and TNF-α at day two, followed by a resolution of cytokine levels by day five, including a ∼7.5-fold reduction in MCP-1, despite increased haemodynamic loading. Endothelial cells retained a cobblestone-like morphology with eNOS undetected, resembling a partially activated phenotype. Compared with steady flow, cardiovascular flow reduced TGF-β1 and IL-8 secretion and decreased FGF-b consumption (∼2.5 fold), suggesting enhanced phenotypic stability.

This study presents the first *in vitro* IA model incorporating a cardiovascular flow waveform and identifies cytokine signatures with potential utility as biomarkers of IA remodelling, highlighting the importance of long-term perfusion for modelling chronic vascular disease.

**Table of Contents Figure:** An *in vitro* model of an intracranial aneurysm was developed to investigate how fluid flow dynamics impact endothelial remodelling and inflammation. Pulsatile cardiac flow promoted stabilisation of inflammatory signalling, which was sustained under a steady flow regime. Cytokine signatures emerged with potential utility as biomarkers of IA remodelling, highlighting the importance of long-term perfusion for modelling chronic vascular disease. The schematic of the cytokine release dynamics used in the graphical abstract below was generated with the assistance of AI-based tools including ChatGPT (v5.5) and M365 Copilot to align with key results from this manuscript.

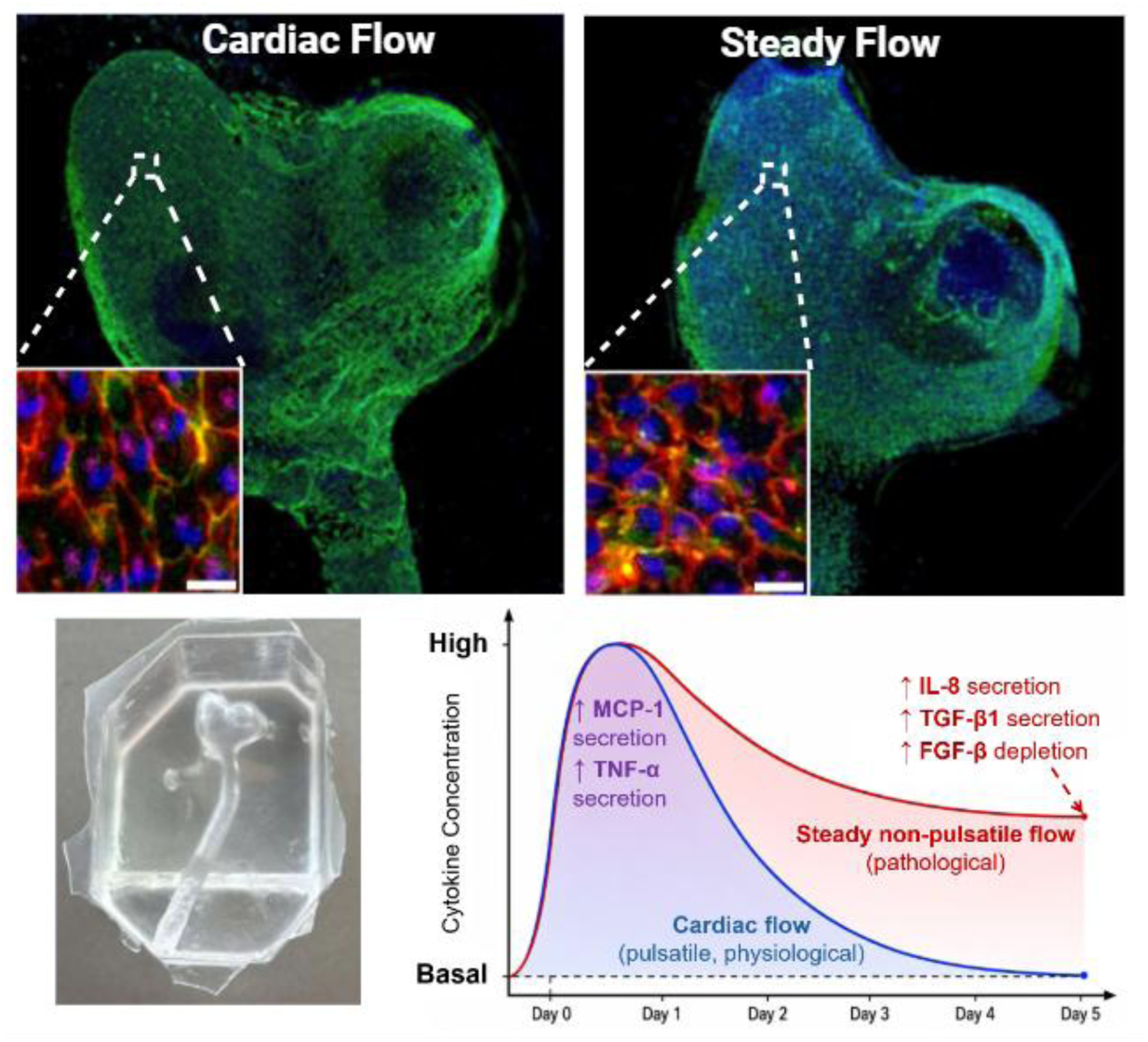

## 1 Introduction

Intracranial aneurysms (IAs) are an outpouching of the intracranial arterial wall that can rupture, resulting in subarachnoid haemorrhage with 30-50% mortality and significant disabilities in survivors.^1–4^ If detected early, clinicians can evaluate and assess the risk of a rupture event against procedural risks to determine the most appropriate patient treatment.^5,6^ To do this, clinicians can refer to simple risk prediction methods that consider morphological characteristics and patient condition.^7,8^ However, these scoring systems have been reported as less informative compared to clinical opinion and professional preference for determining the best treatment strategy.^7,9,10^ To improve the accuracy and reliability of IA rupture prediction models, the underlying mechanisms driving IA growth and rupture should be considered, including changes in haemodynamic stresses, vascular tissue remodelling and molecular signatures, such as alterations in biomarker expression.

Endothelial cells (ECs) lining the lumen of blood vessels play a critical role in maintaining vascular homeostasis in response to mechanical forces exerted by blood flow.^11^ These forces include wall shear stress (WSS), cyclic stretch due to vessel dilation and contraction, and blood pressure.^12,13^ Additionally, the complex arterial architecture in the body can create heterogeneous blood flow patterns that give rise to distinct EC phenotypes and functions.^11^ For example, it has been well established in the literature that ECs exposed to unidirectional, high shear, laminar flow regimes exhibit a quiescent endothelial phenotype.^11,12,14^ Comparatively, where disturbed, non-uniform or reciprocating flow can be present at arterial bifurcations and regions of high curvature, ECs can exhibit a pathogenic phenotype associated with vascular disease.^12,14,15^ These phenotypic changes can be reflected in morphological changes such as cell shape and alignment, expression of surface markers, and the release of soluble factors such as cytokines and chemokines into the surrounding environment.^14,16–22^

IA growth and remodelling is a complex process, thought to be underpinned by geometry and subsequent haemodynamic and biomechanical forces that drive cellular changes.^23^ However, the EC phenotype and remodelling mechanisms involved remain poorly understood and are instead largely hypothesised based on well-characterised mechanisms implicated in atherosclerosis and other vascular diseases.^23^ Furthermore, published literature remains inconclusive as to whether low or high WSS causes IA rupture. Acknowledging that both low and high WSS can drive aneurysmal change, Meng et al. (2014) proposed two flow-dependent remodelling pathways in IAs: inflammation-mediated wall remodelling with wall thickening caused by low and stagnant flow, and high-WSS driven mural cell mediated wall remodelling with wall thinning. However, this has not yet been validated. Understanding these interactions is essential for developing reliable and objective guidance for physicians on aneurysm rupture risk.

To study these interactions, human IA tissue has been collected for histological studies after autopsy or surgical treatment. However, the variability in location and size of resected tissue, low sample size and heterogeneity of samples limit the conclusions that can be made.^24,25^ Similarly, *in vivo* animal models have been studied, however they often do not reflect human IA biology or dynamics.^23^ While *in silico* models can provide computational fluid dynamics (CFD) analysis and enable predictive modelling of IAs, they too lack consideration for the underlying cellular remodelling mechanisms. Comparatively, *in vitro* bioengineered vascular models can bridge this gap by enabling controlled studies on biological remodelling in relation to fluid dynamics and IA morphology, facilitating new insights into EC dysfunction and the growth and rupture dynamics of IAs.^26^

Various *in vitro* systems have been implemented to study the effect of WSS on ECs, including parallel-plate flow chambers, microfluidic devices and bioengineered IA models.^19,27–33^ These studies generally align with the literature on EC phenotype, showing alignment of vascular ECs with the direction of flow, disordered morphologies within aneurysm domes, and upregulation of inflammatory markers under low or disturbed flow. However, these studies primarily use HUVECs which while robust, can exhibit differential responses to arterial ECs under shear^34,35^ Furthermore, perfusion studies extending beyond 24 hours are rarely achieved, which is essential for understanding the effects of sustained variations in the haemodynamic environment that contribute to aneurysm progression. Levitt et al. (2019) even indicated cell detachment was common in their bioengineered PDMS IA model when exposed to flow for more than 24 hours.^27^ While many fluidic studies have investigated EC phenotype in parallel plate flow chambers and microfluidic devices, these platforms are typically flat, rectangular chambers at the micro-scale that do not fully capture the physiological vessel curvature, lumen geometry or associated flow regimes present *in vivo.*^31,36,37^ In contrast, fluidic studies in cylindrical channels of physiologically relevant dimensions are scarce. Similarly, despite ECs being sensitive not only to the magnitude of WSS but also the temporal and directional variability of the flow regime, *in vitro* analyses of EC phenotype under a physiological cardiovascular flow waveform are limited.^12,14,38–44^ Incorporating physiological flow waveforms into *in vitro* IA models is essential to accurately investigate endothelial behaviour in the context of IAs.

As EC phenotype can be influenced by substrate stiffness, curvature, cell origin and flow regime, we benchmark EC phenotype in large 3D PDMS cylindrical channels (2.2 mm diameter) at steady physiological flow rates for translation to *in vitro* IA models.^12^ We hypothesised that human aortic endothelial cells (HAECs) exposed to well-defined, steady physiological shear stress within cylindrical PDMS channels for over 24 hours (five days total) would adopt a shear-adapted phenotype characterised by a flow-aligned morphology, increased eNOS expression, and minimal inflammatory activation.

Furthermore, while unidirectional pulsatile and cardiovascular flow regimes can elicit a protective signalling response in ECs, the complex geometry of IAs can complicate this relationship.^12,14,41–44^ It is anticipated that perfusion of *in vitro* IA models with a cardiovascular flow waveform will create a more disturbed and oscillatory flow regime within the aneurysm dome compared to steady flow conditions. We therefore hypothesise that HAECs cultured under these conditions will exhibit a more dysfunctional endothelial phenotype compared to steady flow conditions, characterised by a cobblestone-like morphology and enhanced inflammatory and remodelling signalling.

Given that endothelial responses to haemodynamic forces involve both structural adaptation and active cytokine signalling, assessment of EC phenotype requires evaluation of secreted inflammatory and remodelling markers in addition to morphological imaging. We therefore characterise our system through immunofluorescence imaging of immortalised HAECs with analysis of secreted factors under steady non-pulsatile and cardiovascular flow conditions. To our knowledge, we present the first *in vitro* IA model system that incorporates a physiologically relevant cardiovascular flow waveform and captures the temporal endothelial response after two and five days.

## 2 Results

### 2.1 Engineering idealised and personalised models of IA haemodynamics

Using STL files extracted from microcomputed tomography (µCT) imaging of the straight channels (**Figure 1**b), CFD analyses enabled the evaluation of WSS and velocity profiles. Simulation results indicated average WSS values of 0.30 ± 0.11 Pa under low flow (31 mL/min), 0.87 ± 0.37 Pa at medium flow (75 mL/min) and 1.60 ± 1.11 Pa at the high flow condition (121 mL/min) (Figure 1c), all within the physiological range reported across the Circle of Willis.^45,46^ However, high flow conditions created a heterogeneous WSS profile along the length of the cellularised region, with localised pockets of high (>1.8 Pa) and low (<0.5 Pa) WSS. As the flow exited the barbed connector and entered the cellularised region, the flow path and velocity profile was disturbed, stabilising approximately 17 mm into the channel across all conditions (Figure 1d, e-g).

**Figure 1:**
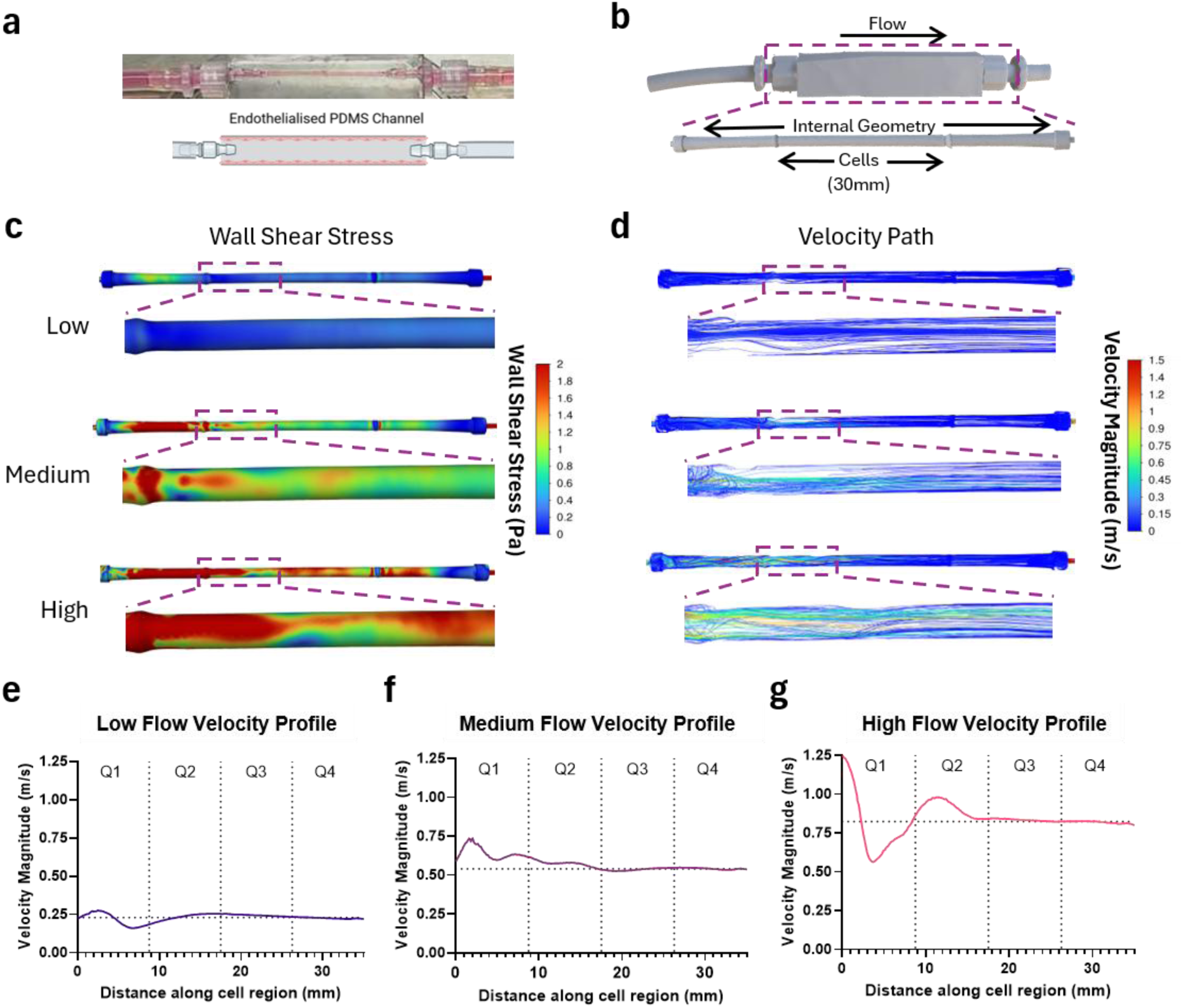
Computational fluid dynamics analysis through straight PDMS channels at Low (31 mL/min), Medium (75 mL/min) and High (121 mL/min) flow conditions. (a) Image of straight PDMS channel connected to tubing and filled with culture media (pink). Graphic depicts cell location, created with BioRender.com. (b) STL of the PDMS channel and internal geometry. (c) Wall shear stress distributions through the internal geometry. (d) Velocity path distributions through the internal geometry. Centreline velocity profile at (e) low, (f) medium and (g) high flow across cellularised region. Horizontal dotted lines in figures e - g represent the average velocity across 20 mm to 35 mm of the cellularised region. Vertical dotted lines represent quartile segments the PDMS channels were sectioned into.

IA modes were fabricated with an accuracy of 85.2 ± 1.7% compared to the original segmented model, with negligible expansion under pressure (**Figure 2**a-c, FigureS4). CFD simulations with culture media revealed *in vitro* WSS values ranging from 0.94 ± 0.66 to 3.14 ± 2.45 Pa under the cardiovascular flow regime that overlapped with the steady-flow regime (1.47 ± 1.03 Pa), and the simulated physiological blood flow range (2.12 ± 1.60 to 4.34 ± 3.20 Pa, Figure 2g and h). Spatial analysis of WSS across perfused IA models are provided in Figure S5.

**Figure 2:**
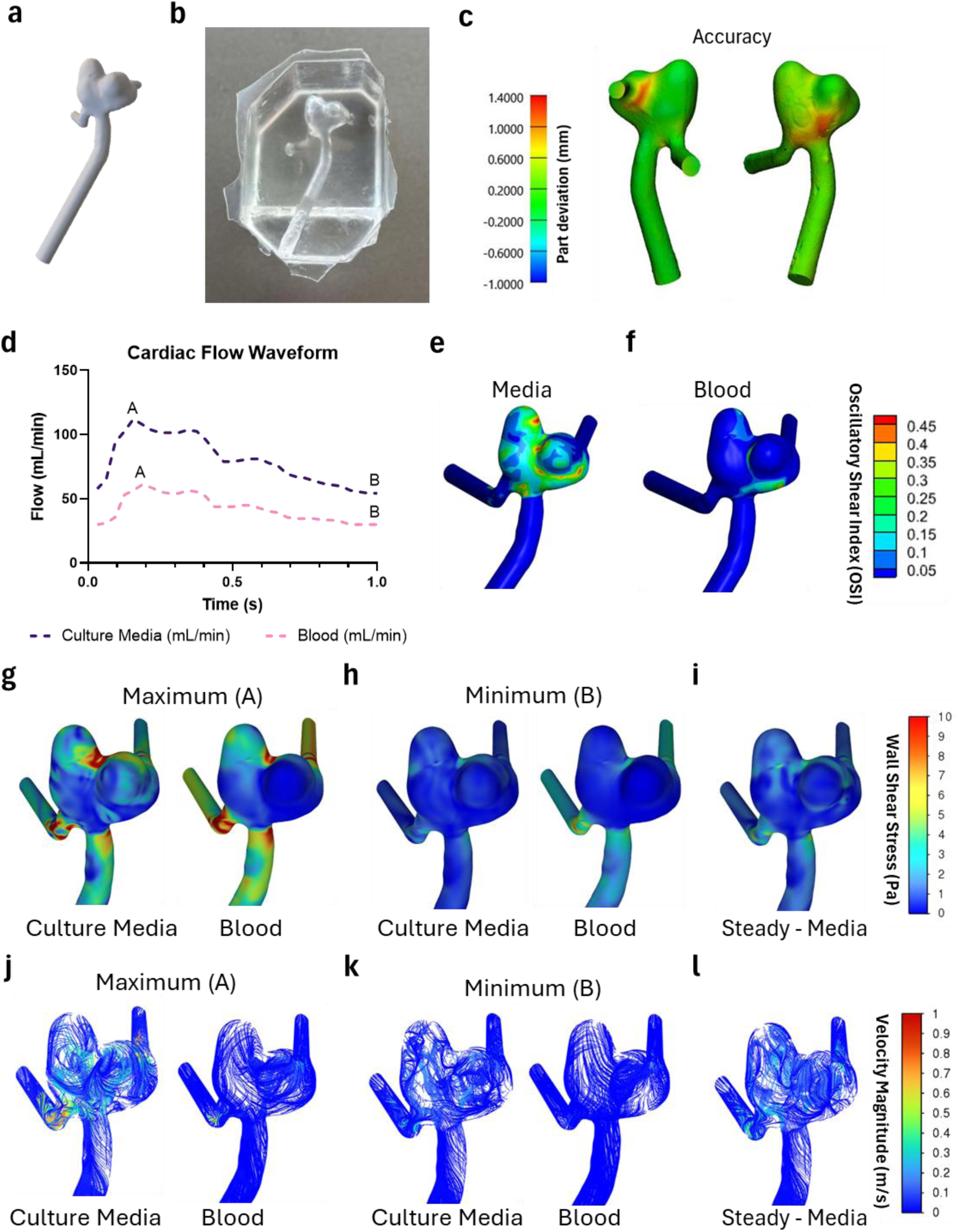
**Computational Fluid Dynamics (CFD) analysis of blood and culture media flow through a bi-lobed middle cerebral artery aneurysm model**. (a) STL of bi-lobed aneurysm selected for modelling. (b) *In vitro* PDMS model of intracranial aneurysm. (c) Geometric accuracy of *in vitro* PDMS aneurysm model. (d) Cardiac flow waveforms analysed comparing physiological blood flow with experimental culture media conditions. Maximum and minimum flow points denoted ‘A’ and ‘B’ respectively. Oscillatory shear index (OSI) profiles of (e) culture media and (f) blood under a cardiac waveform. Wall shear stress profiles of blood and culture media analysed at the (g) maximum and (h) minimum point of the cardiac waveform. (i) Wall shear stress profile of culture media at steady flow of 60mL/min. Velocity profiles of blood and culture media analysed at the (j) maximum and (k) minimum point of the cardiac waveform. (l) Velocity profile of culture media at steady flow of 60mL/min.

Although, achieving these WSS profiles required an increased media flowrate to compensate for the lower viscosity of media relative to blood (Figure 2d). This resulted in an accelerated velocity profile (Figure 2j and k) and generated a more disturbed, less streamlined flow pattern compared to physiological blood flow. Consequently, elevated and more spatially variable oscillatory shear index (OSI) values were observed within the aneurysm dome under the cardiovascular flow regime (Figure 2b).

### 2.2 Endothelial monolayer density is impacted by flow and geometry

High cell coverage was retained across the entire length of the PDMS channels for all flow conditions, as depicted in stitched fluorescent images spanning Q2-Q4 (**Figure 3**a). HAECs were seeded at 120,000 cells/cm^2^, increasing to an average of 208,480 ± 11,306 cells/cm^2^ under static conditions (Figure 3b). At low flow (31 mL/min), cell density remained above the initial seeding density across the entire channel (>136,000 cells/cm^2^). Under medium flow (75 mL/min), a significant reduction in cell density was observed in Q1 (81,623 ± 21,985cells/cm^2^) that recovered downstream in Q2-Q4 to an average of 158,698 ± 52,179 cells/cm^2^. At the highest flow (121 mL/min), cell density was significantly lower across the entire channel relative to static conditions (116,190 ± 12,165 cells/cm^2^, P < 0.01). Under all conditions, cell viability in the first quarter of the channel remained at or above 90% (Figure 3c). Figure S6 depicts the IF results of the viability (dead) and IL-6 staining, where IL-6 was not detected.

**Figure 3:**
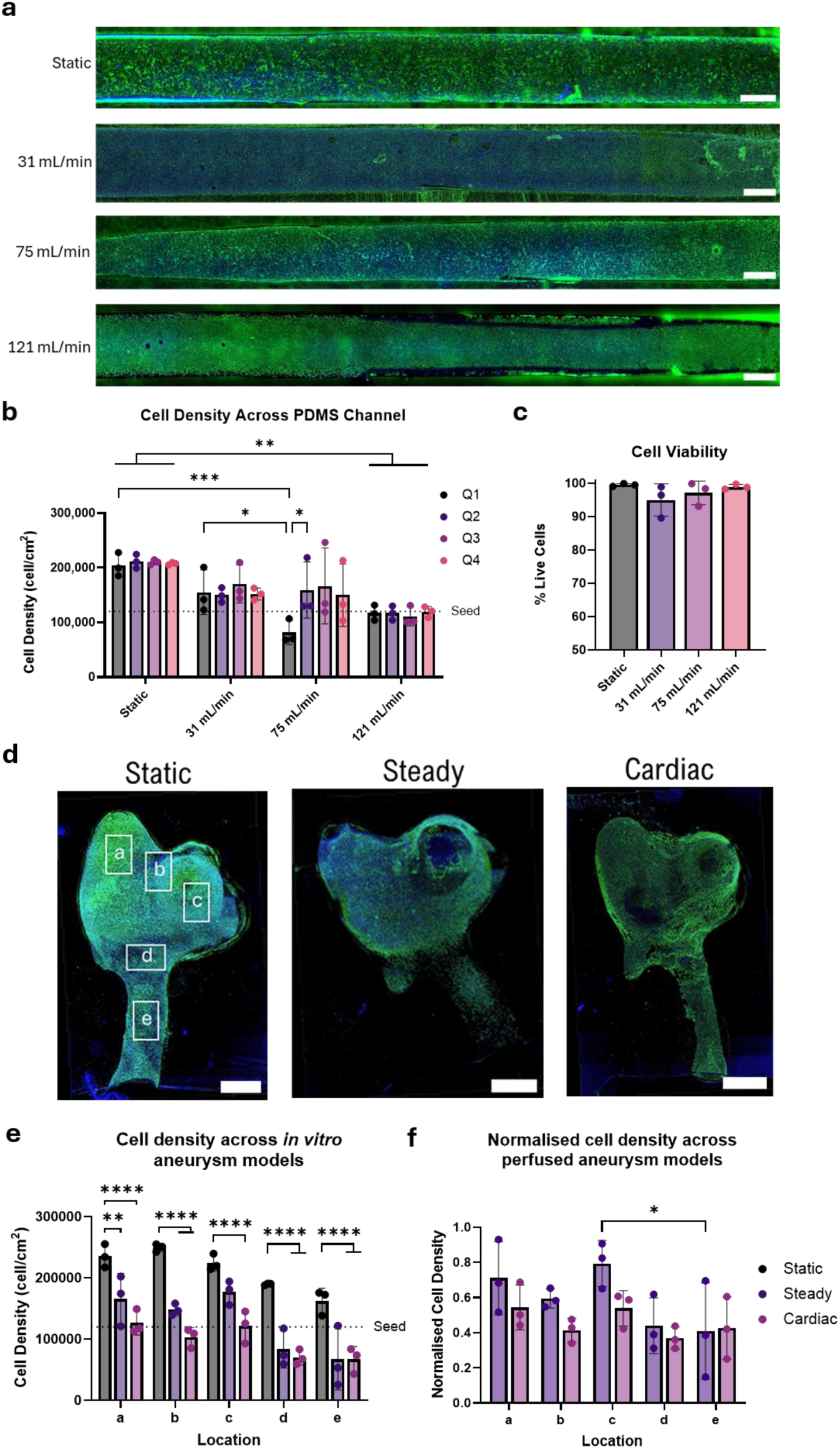
**Human aortic endothelial cell (HAEC) density and viability on polydimethylsiloxane (PDMS) channels and intracranial aneurysm (IA) models after five days of perfusion**. (a) Stitched fluorescent images of HAECs fluorescing green (GFP) and stained for DAPI (blue) between Q2 and Q4 on PDMS channels. Scale represents 1 mm. (b) Cell density across Q1 to Q4 of PDMS channels. (c) Viability of HAECs determined by staining for dead cells using Live-or-Dye^TM^ 568/583 viability stain (1:1000, Cat. No. 32005, Gene Target Solutions) on Q1 of PDMS channels. (d) Stitched fluorescent images of HAECs fluorescing green (GFP) and stained for DAPI (blue) and F-actin (green) on IA models after perfusion. Scale bar represents 2 mm. (e) Quantification of cell density across five locations of IA models. (c) Normalised cell density of perfused IA models against static cultures. Due to the large size of the IA models and straight channels, minor imaging artefacts related to image stitching, fluorescent shading, blending, surface defects, surface depth variability and narrowing or widening of the channel from sectioning techniques can be observed.

In contrast, perfused PDMS IA models exhibited regional variability in endothelial coverage. While high cell coverage was also retained across the aneurysm dome, cell detachment was pronounced at the aneurysm neck and inlet regions under flow (Figure 3d, Figure S7). To quantify spatial differences, each cellularised IA model was imaged across five distinct locations (Figure 3d Static) and cell density quantified (Figure 3e). Two-way ANOVA revealed a significant main effect of both location (P<0.0001) and flow (P<0.0001) on cell density. No significant interaction between location and flow was detected (P = 0.2537), indicating that the influence of flow on cell density was consistent across regions. The spatial variation in cell density under static conditions independent of flow suggests local geometric effects on seeding variability. Normalisation of cell density to static samples revealed spatial differences only under steady flow between the right lobe (location ‘c’) and the inlet (location ‘e’) (P = 0.0402), coinciding with the lowest and highest WSS regions respectively (Figure 3f, Figure S5).

Notably, high cell density was maintained in the left lobe (location ‘a’) under flow despite the elevated regional WSS (Figure S5) comparable to the aneurysm inlet (locations ‘d’ and ‘e’). Consistent with this observation, analysis across all models demonstrated no relationship between local cell density and WSS magnitude (Figure S8).

### 2.3 Endothelial morphology is preserved under flow, with localised changes in junctional organisation and alignment

CD31 and ZO-1 antigens were co-stained in the straight channel models to evaluate junctional integrity under increasing flow (**Figure 4**a and Figure S9). CD31 expression remained high across all conditions, quantified by mean fluorescence intensity (MFI) (Figure 4d). In contrast, ZO-1 co-localisation with CD31 was significantly reduced at 121 mL/min compared with static conditions and 75 mL/min flow (Figure 4b, P = 0.0275 and P = 0.005 respectively). In perfused IA models, CD31 MFI was significantly reduced at locations ‘b’ and ‘e’, corresponding to regions of higher WSS (Figure S5) at the intersection of the bi-lobed aneurysm dome and the inlet. Isotype staining is presented in Figure S9.

**Figure 4:**
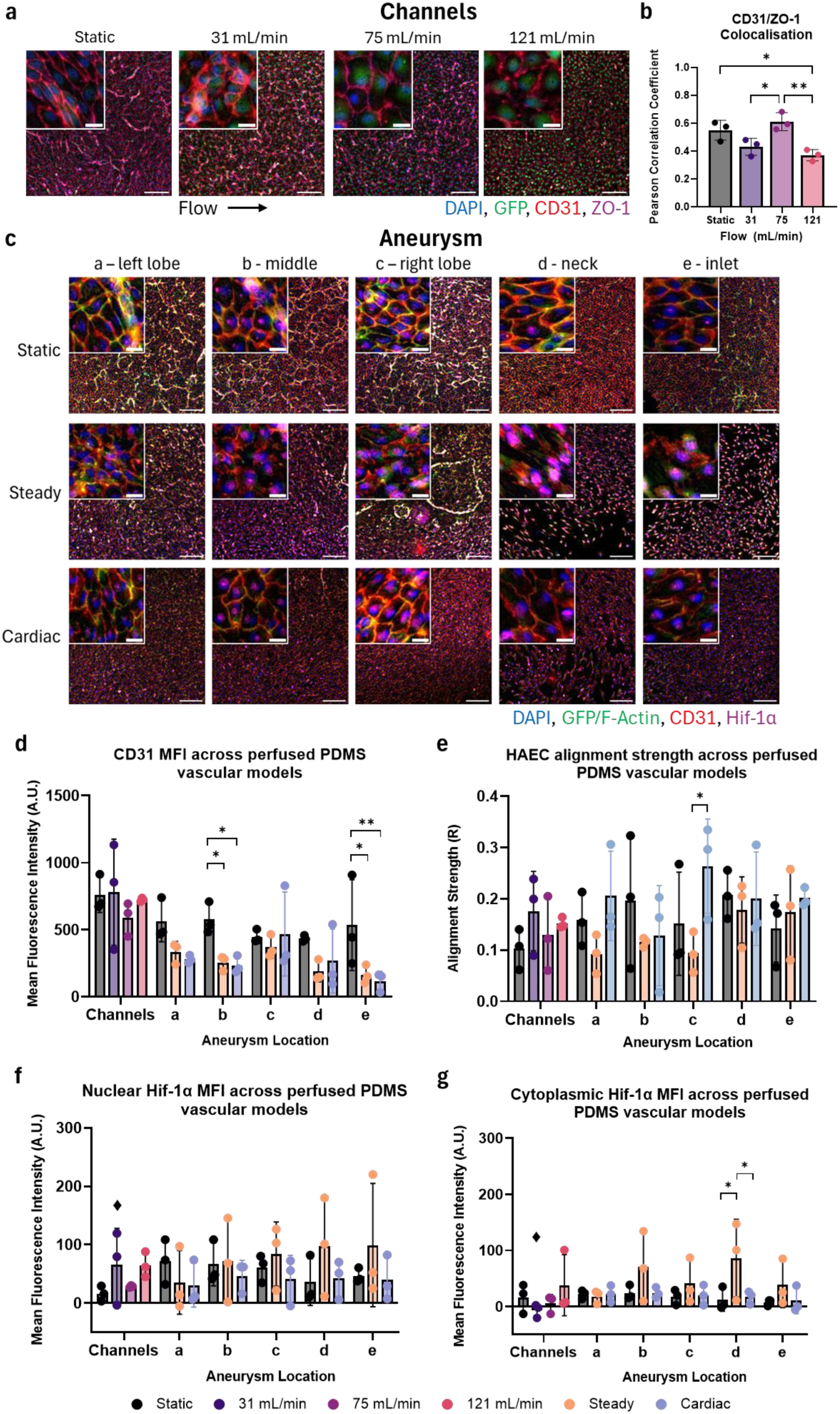
**Morphology and alignment of human aortic endothelial cells (HAECs) under flow in straight polydimethylsiloxane (PDMS) vascular models and intracranial aneurysm (IA) models**. (a) Immunofluorescent images of GFP (green) fluorescing HAECs on PDMS channels stained for DAPI (blue), CD31 (red) and ZO-1 (magenta). Scale of embedded image represents 20 µm. Scale of larger low-resolution image represents 200 µm. (b) Pearson correlation coefficient of CD31 and ZO-1 using the Colocalization plugin of Fiji. (c) Immunofluorescent images of HAECs on IA models stained for F-actin (green), DAPI (blue), CD31 (red) and Hif-1α (magenta). Scale of embedded image representtos 20 µm. Scale of larger low-resolution image represents 200 µm. (d) Mean fluorescence intensity (MFI) of CD31. (e) Alignment strength of HAEC CD31 patterns. (f) MFI of Hif-1α localised to the nucleus. (g) MFI of Hif-1α localised to the cytoplasm.

Cells exhibited a predominantly cobblestone-like morphology across all straight channel and aneurysm models under perfusion, which was apparent at day one of 0.2 mL/min flow (Figure S10). This contrasted with the elongated and randomly oriented morphology observed across static cultures (Figure 4a and c). Alignment analysis using the OrientationJ plugin of Fiji on CD31-stained images revealed weak alignment strength in straight channels (R < 0.3, Figure 4e) with a minor preference for cells to orient perpendicular to flow (<10% of pixels, Figure S9). Similarly, weak alignment was observed in perfused IA models (R < 0.4, Figure 4e) with greater alignment observed under cardiac flow in the right aneurysm lobe (location ‘c’) compared to steady flow conditions (Figure 4c and e, P = 0.0271, Figure S11). At this location, the velocity profile at peak systole of the cardiac cycle shows a more pronounced rotational pattern and formation of a small vortex (Figure 2j – culture media) that was absent under steady flow (Figure 2l).

Despite a lack of quantitative significance, some notable alignment patterns were observed across perfused models. For example, more pronounced cell alignment was seen in regions of lower cell density, particularly at aneurysm inlet locations ‘d’ and ‘e’. Figure S12 depicts a clear transition between a cobblestone to an aligned morphology as cells become sparse and F-actin staining more diffuse. However, it is unknown if this alignment is parallel or perpendicular to flow due to the heterogeneity in the velocity flow path across the aneurysm dome and throughout the cardiac cycle. Meanwhile, in regions where surface grooves from 3D-printed layer lines were present, cells showed no preferential alignment with these grooves, as depicted in Figure S13.

No significant differences were observed in Hif-1α expression with increasing flow across straight channels. However, increased nuclear localisation was evident at 31 mL/min relative to static conditions. Similarly, limited differences were observed across perfused IA models, with elevated cytoplasmic expression observed under steady flow at the aneurysm neck. Isotype staining is presented in Figure S14.

HAECs continued to proliferate under both static and flow conditions in straight channel experiments, with 2 to 15% of the cell population expressing Ki67 across groups (**Figure 5**a and b). eNOS expression, a key indicator of cell phenotype, was comparable to isotype controls for many samples across conditions (Figure 5c). Isotype staining is presented in Figure S15.

**Figure 5:**
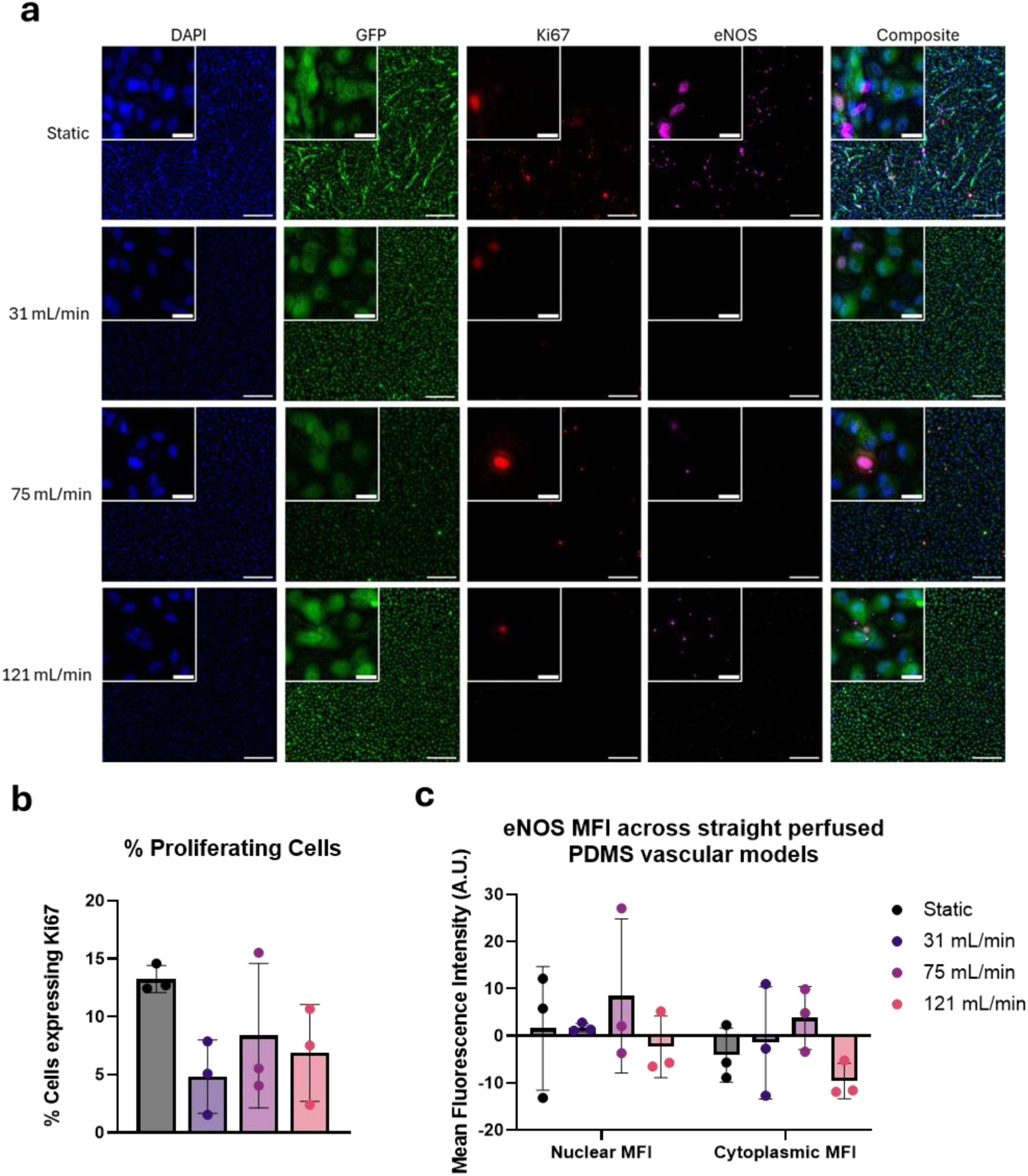
**Ki67 and eNOS expression in human aortic endothelial cells (HAECs) under static and flow conditions in straight vascular models**. (a) immunofluorescent images of GFP (green) fluorescing HAECs on PDMS channels stained for DAPI (blue), Ki67 (red) and eNOS (magenta). Scale of embedded image represents 20 µm. Scale of larger low-resolution image represents 200 µm. (b) Percentage of cells expressing Ki67. (c) Mean fluorescence intensity (MFI) of eNOS-stained samples corrected against isotype controls. Negative results indicate signal was undetectable.

### 2.4 Endothelial growth factor uptake and release increases with flow

Of the 13 analytes assessed using the 13-plex LEGENDplex^TM^ Human Growth Factor Panel, seven were reliably detected within perfused media reservoirs. Five analytes (TGF- β1, FGF-B, VEGF, Angiopoietin-2, and SCF) demonstrated flow-dependent trends and are presented in **Figure 6**, while PDGF-AA and EGF results are provided in Figure S16. Original measurements are presented in Figure S17 and S18 with the volume-corrected and cell-normalised analyte rates of change presented in Figure 6.

**Figure 6:**
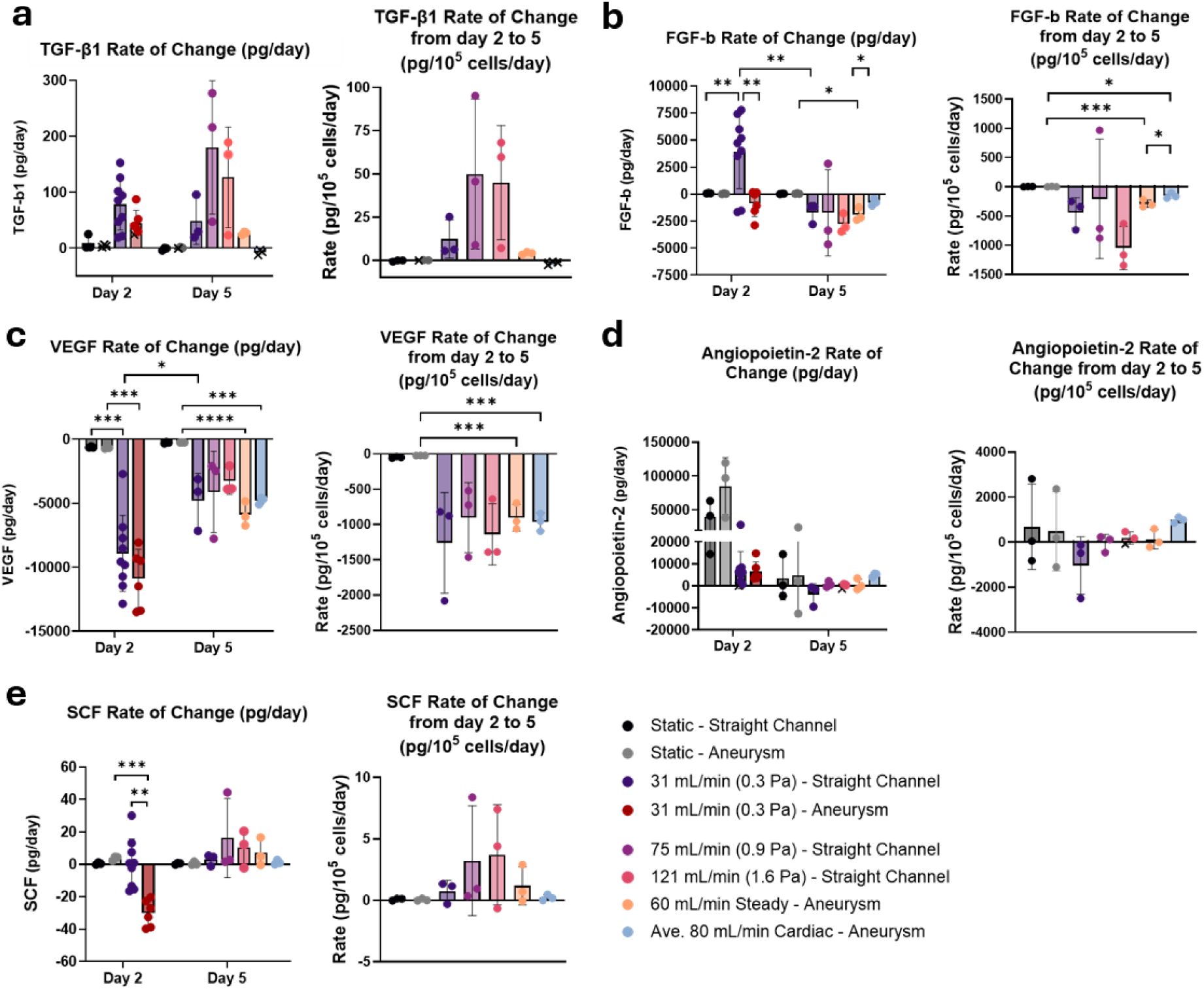
**LEGENDplex^TM^ quantification of endothelial growth factors at day 2 and 5 of perfusion of PDMS vascular models**. (a-e) Volume-corrected and cell density-normalised rate of change of (a) Transforming growth factor-β1 (TGF-β1), (b) Fibroblast growth factor-basic (FGF-b), (c) Vascular endothelial growth factor (VEGF), (d) Angiopoietin-2 and (e) SCF. Measurements below the limit of detection were considered undetected and have been marked with an ‘x’. Cell density normalisation based on quantification of cell density in Figure 3.

Despite frequent measurements below the LOQ, TGF-β1 displayed qualitative differences in detection pattern and calculated secretion rates. In straight channel models, TGF-β1 secretion increased under flow at day two compared to static conditions and remained elevated under steady flow conditions at day five (Figure 6a). In contrast, in IA models TGF-β1 was only reliably detected at day five under steady flow. Notably, this condition utilised the largest media volume (∼36 mL compared to 16 mL under cardiovascular flow), indicating detectable TGF-β1 accumulation despite greater dilution.

In straight channel perfusion cultures, FGF-b secretion was significantly increased at day two compared to static conditions (P = 0.0099), indicated by a positive rate of change. By day five, rates became negative across all flow conditions indicating net consumption. In IA models, FGF-b consumption was approximately 2.5 fold higher under steady flow compared to cardiovascular flow at day five (-1922 ± 609.6 pg/day vs - 763.6 ± 293.2 pg/day, P = 0.0255).

Due to reduced media volume in static cultures (∼1 mL), limited VEGF was available for continued cellular consumption (Figure S17 and S18). As a result, VEGF consumption was significantly increased under flow in straight channels, with a modest reduction by day five of exposure at 31 mL/min (P = 0.0471). In IA models, VEGF consumption remained significantly upregulated under both steady and cardiovascular flow conditions at day 5 relative to static cultures.

Angiopoietin-2 secretion rate was non-significantly elevated under static conditions at day two compared to flow. While no significant differences were observed at day 5, cell-normalised secretion was slightly elevated under cardiovascular flow compared to steady flow conditions Only FGF-b rate of change (Figure 6b) and SCF rate of change (Figure 6e) were significantly reduced in the IA models at day two compared to straight channels. At this time point, both models were exposed to identical flow rates and exposure duration, indicating an effect of geometry and subsequent flow pattern on early FGF-b and SCF response. SCF was not added as an exogenous growth factor, and therefore trace levels are likely a result from supplemented FBS.

### 2.5 Long-term perfusion uncovers stable cytokine release profiles of endothelial inflammation

Of the 13 analytes assessed using the 13-plex LEGENDplex^TM^ Human Inflammation Panel, four were reliably detected including MCP-1, TNF-α, IL-6 and IL-8. Raw analyte measurements are provided in Figure S17 and S18, with the volume-corrected and cell-normalised analyte rates of change presented in **Figure 7**.

**Figure 7:**
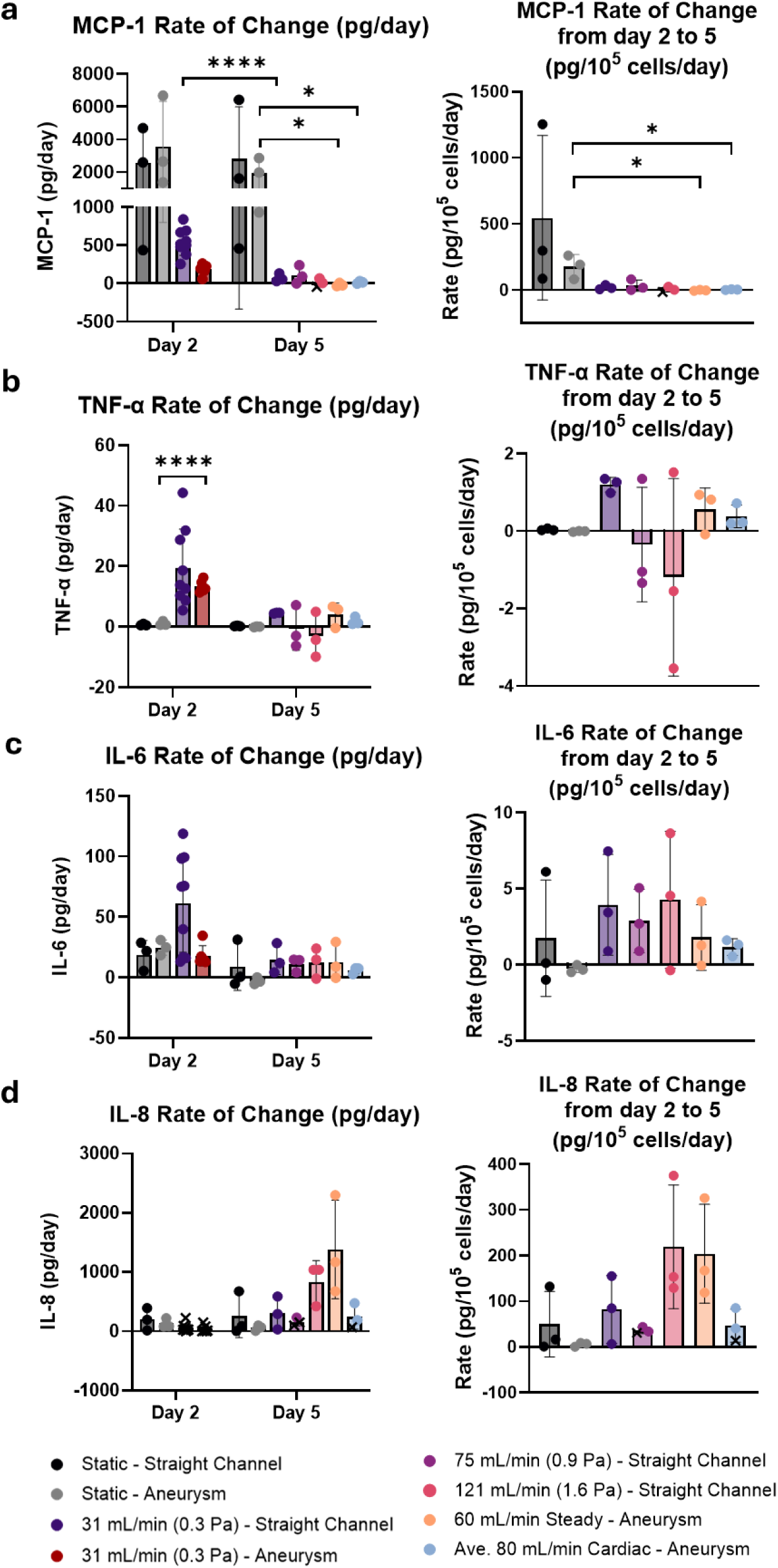
**LEGENDplex^TM^ quantification of endothelial inflammatory cytokines at day 2 and 5**. (a-d) volume-corrected and cell density-normalised rate of change of (a) Monocyte chemoattractant protein-1 (MCP-1), (b) Tumour Necrosis Factor-alpha (TNF-α), (c) Interleukin-6 (IL-6), (d) Interleukin-8 (IL-8). Measurements below the limit of detection were considered undetected and have been marked with an ‘x’. Cell density normalisation based on quantification of cell density in Figure 3.

MCP-1, TNF-α and IL-6 secretion rates (Figure 7a-c) were upregulated under multiple flow conditions at day two and stabilised by day five despite increasing WSS magnitude. In straight channel models where 31 mL/min was maintained beyond day two, MCP-1 secretion rate decreased 7.5-fold by day five (543.5 pg/day vs 72.18 pg/day, P<0.0001). Although TNF-α secretion rate was significantly elevated in IA models at day two compared to static cultures (13.29 ± 1.85 pg/day vs 1.08 ± 0.74 pg/day, P < 0.0001), secretion rates were comparable across all flow conditions by day five. Elevated IL-6 secretion was also observed under flow at day two in straight vascular models, which decreased by day five.

In contrast, IL-8 (Figure 7d) was not detected under flow at day two in either straight channel or IA models, while elevated levels were observed in static cultures. By day five, IL-8 was detected under steady flow conditions in straight channels and 60 mL/min in IA models, indicating continued or upregulated IL-8 secretion. IL-8 secretion rate also appeared to be reduced in IA models under cardiovascular flow compared to steady flow at day five.

## 3 Discussion

Vascular remodelling mechanisms driving the growth and rupture of unstable IAs remain poorly understood, limiting the clinical utility of existing rupture prediction models. To address this, the present study investigated endothelial phenotypic responses to steady and cardiovascular flow patterns within *in vitro* idealised and patient-specific IA models.

Across straight cylindrical channels similar in size to the middle cerebral artery (2.2 mm diameter), high cell density (> 116,190 cells/cm^2^) and viability (> 90%) were maintained over five days of perfusion, achieving physiological flow rates up to 1.6 Pa shear stress (121 mL/min). Flow initiation triggered an acute inflammatory and remodelling response, characterised by flow-dependent changes in TNF-α, IL-6, MCP-1, TGF-β1, FGF-b and VEGF over the first two days of perfusion. This was followed by stabilisation towards baseline across day 2 to 5 despite increasing shear stress magnitude. These findings suggest endothelial adaptation to sustained haemodynamic stress and highlights the importance of extending *in vitro* haemodynamic studies beyond the commonly adopted 24 to 48-hour window. This work represents one of the first successful millimetre-scale, cylindrical PDMS vascular models supporting multi-day perfusion with an endothelialised lumen under physiologically relevant flow conditions. This platform therefore enables the study of sustained vascular responses relevant to chronic vascular disease pathophysiology and progression.

Building up on the complexity and physiological relevance of this platform, patient-specific IA models were endothelialised and exposed to a cardiovascular waveform. To our knowledge, this represents the first *in vitro* study to implement a cardiovascular waveform within a patient-specific IA model, enabling investigation of flow-driven endothelial behaviour under aneurysm-relevant conditions.

It was hypothesised that prolonged exposure to steady physiological shear in idealised channels would induce a quiescent, shear-adapted phenotype marked by a flow-aligned morphology, increased eNOS expression, and minimal inflammatory activation.^14,16–22^ However, this phenotype was not observed in our studies. Instead, HAECs exhibited a cobblestone morphology, unchanged eNOS expression across shear conditions, and an acute inflammatory response that stabilised over time. Where cell alignment with flow and eNOS upregulation are hallmarks of a quiescent phenotype, their absence may reflect partial activation towards a more dysfunctional phenotype in this system.^14,16–19^ Similar findings were observed in the IA models, where alignment was restricted to regions of elevated flow or low cell density such as the aneurysm neck. Reduced CD31 expression at the inlet and intersection of the aneurysm lobes, with elevated Hif-1α at the aneurysm neck under steady flow further suggest that impinging and spatially variable flow, rather than WSS magnitude alone, could modulate EC phenotype within IA geometries.

In contrast to imaging-based markers, cytokine and biomarker profiling can be a highly informative, non-invasive method for evaluating EC phenotype and dysfunction both experimentally and clinically in relation to IA rupture risk.^20,21^ Here, the LEGENDplex^TM^ multiplex cytokine analysis revealed significantly elevated FGF-b consumption with non-significant elevation in TGF-β1 and IL-8 secretion under steady flow compared to cardiovascular flow. These expression patterns could indicate a more metabolically active, pro-inflammatory and adaptive phenotype under steady flow conditions, characterised by enhanced growth factor consumption, cytokine secretion and remodelling-associated signalling.^47^ Given the role of TGF-β1 and FGF-b in ECM synthesis and intimal wall thickening, and IL-8 in the recruitment of neutrophils and EC permeability, the slower steady flow profile could create regions of low flow and stagnation in the aneurysm dome that lean into the inflammation-mediated wall remodelling hypothesis without complete pathological dysfunction.^23,48–56^ In contrast, cardiovascular flow was associated with reduced inflammatory output and growth factor demand, consistent with a more shear-adapted and atheroprotective phenotype.^12^ Cardiovascular flow could therefore accelerate phenotypic adaptation and reduce susceptibility to a pathological state, consistent with previous studies using parallel-plate flow and cone and plate devices, despite the complex IA geometry producing regions of high OSI ^43,44,48^.

In previous *in vitro* and *in vivo* studies, abnormal TGF-β1 signalling has been implicated in aortic aneurysm wall integrity, FGF-b signalling in aneurysmal fibrosis and subarachnoid haemorrhage, and IL-8 has been upregulated in plasma collected from patients with intracranial aneurysms. ^56–60^ Additionally, TNF-α, IL-6, MCP-1 and IL-1β have also been detected in the serum of patients with aneurysms and have been associated with rupture ^60–63^. Collectively, these findings posit that profiling EC cytokine release can provide insight into the degree of endothelial dysfunction and remodelling activity in IAs, with potential to complement clinical rupture-risk prediction frameworks.

Importantly, the *in vitro* models developed in this study are not intended for direct patient-specific rupture risk assessment on a case-by-case basis. Rather, they provide a control experimental platform to systematically characterise endothelial responses to defined haemodynamic and biochemical stimuli. These models can therefore be leveraged to inform and refine existing rupture-risk prediction frameworks such as the PHASES and UIATS scores, for example through the integration of mechanistic biomarkers into statistical or machine learning-based predictive models.^7,8,64,65^ By enabling controlled investigation of relationships between flow conditions, endothelial phenotype and biomarker expression, this approach may support the development of more biologically informed risk stratification tools. However, given the complexity of the underlying signalling networks involved, including the context-dependent and synergistic behaviour of cytokines and growth factors, interpretation of biomarker panels remains challenging. Further investigation using controlled *in vitro* vascular models alongside clinical validation is required to determine how these molecular signatures can be robustly integrated into rupture risk prediction.^20,21^

While the emergence of the observed phenotype in response to steady physiological flow regimes is inconsistent with the general literature, idealised studies characterising EC phenotype in millimetre-scale PDMS cylindrical channels under physiological flow are scarce.^31^ This raises the possibility of differences in the mechanotransduction phenotype of telomerase-immortalised HAECs, alongside geometry- and system-specific effects such as pressure, flow regime and endothelial cell matrix (ECM) coating.

Due to the nature of peristaltic pumps, pressure and flow were inherently coupled in our system, with pressure increasing with flowrate. Although pressure is rarely reported in fluidic studies despite its close relationship with flow, recent *in vitro* studies have shown that elevated pressure can cause reduced eNOS expression and induce perpendicular cellular alignment to flow even under physiological shear.^66–68^ Therefore, pressure may have contributed to the effects observed here. Furthermore, despite applying a steady flow regime through the idealised cylindrical channels, a variable velocity flow path was created upon entry into the PDMS channels. The high flow velocity, model curvature and geometric scale therefore likely influenced the resulting flow regime, where disturbed flow has been strongly associated with endothelial dysfunction related to a cobblestone morphology and low eNOS expression.^69,70^ While common *in vitro* fluidic systems such as parallel-plate flow chambers and microfluidic devices can impose controlled, uniform flow fields, they lack the three-dimensional curvature and continuous circumferential lumen relevant to blood vessels which can influence alignment and mechanotransduction.^37,71^ Most evidence for flow-mediated endothelial alignment and eNOS upregulation is derived from these simplified systems, even though endothelial phenotype varies considerably *in vivo* with the local haemodynamic environment.^11^ Therefore, the responses observed in millimetre-scale curved PDMS vascular models may reflect adaptation to complex flow conditions rather than a failure to reproduce canonical endothelial behaviour observed in flat, uniform-flow systems. Haemodynamic environments can vary considerably between these fluidic platforms despite application of a similar average WSS, emphasising the need for CFD-based characterisation of flow patterns for meaningful interpretation and comparison of endothelial mechanobiology across fluidic platforms.^71,72^ In addition, while fibronectin is beneficial for EC adhesion to PDMS devices, its deposition *in vivo* during vascular injury, inflammation, and under disturbed flow conditions, can promote shear-related proinflammatory signalling.^73–76^ Together, these factors may bias EC phenotype towards partial activation.

While some *in vitro* IA models have previously been created,^19,27,28^ we build on these models by implementing a cardiovascular flow waveform driven by a custom pneumatic pump to better reflect aneurysm haemodynamics. Further, the observed stabilisation of cytokine release despite sustained haemodynamic stress suggest that short-term perfusion studies may capture transient adaptation rather than a steady endothelial phenotype. Levitt et al. (2019) also indicate cell detachment was common after 24 hours of perfusion, further emphasising the reliability of the system developed in this study with sustained cell attachment over five days. These findings highlight the importance of flow waveform and flow duration on endothelial mechanobiology and provides a framework upon which more comprehensive multi-omic, longitudinal, multi-cellular studies can be built to further investigate IA remodelling mechanisms.

Although distinct quiescent and dysfunctional endothelial phenotypes have been widely correlated with flow regime,^12,14,38^ and inflammation-driven and mural-cell mediated remodelling mechanisms have been linked to thick- and thin-walled aneurysmal regions,^23^ the precise EC phenotype and dominant remodelling mechanisms could not be conclusively resolved in this *in vitro* IA model. This limitation likely reflects the inherent complexity of the endothelial mechanotransduction response which is not yet completely understood.^12^ While this study provides a foundation for future development, several experimental limitations restrict the physiological relevance of the model and extent to which these remodelling responses can be characterised.

This model is limited in its similarity to true vascular mechanics and biology. This study only implemented endothelial cells, however vascular and aneurysmal remodelling dynamics are significantly impacted by smooth muscle cells and fibroblasts as well as other structural extracellular matrix components such as the internal elastic lamina.^77^ Future *in vitro* IA models should aim to incorporate these components to enhance physiological relevance of the model for investigating complete aneurysm remodelling dynamics. In addition, telomerase-immortalised HAECs were used in this model due to poor growth characteristics and lack of CD31 expression in human brain endothelial cells (HBEC-5i), which limited their suitability for extended culture and perfusion. While this improved experimental robustness, it reduced the physiological relevance of the model to cerebral vasculature. Alternatively, patient-specific endothelial cells isolated from peripheral blood, such as blood outgrowth endothelial cells (BOECs), peripheral blood-derived endothelial colony forming cells (PB-ECFCs), or induced pluripotent stem cell-derived endothelial cells (iPSC-ECs) could integrate patient-specific biology alongside patient-specific structure and mechanics.^78^

Despite providing more comprehensive imaging and secretome analyses than existing *in vitro* IA models,^19,27^ incorporating broader analysis and multi-omic approaches would further strengthen interpretations of mechanotransduction and remodelling processes. This could include gene expression analysis like Levitt et al. (2019) and metabolomics analysis where some lipids have been associated with aneurysm rupture *in vivo.*^27,62^ Additionally, EC phenotype comparisons between the two lobes of the aneurysm were only evaluated using a single IF panel, while LEGENDplex^TM^ cytokine readouts were not spatially resolved and instead provided a holistic measure of system response. As such, direct correlation between these modalities is limited in the IA models. In contrast, analysis in idealised channels enables controlled isolation of flow conditions, allowing direct correlation between defined shear stress profiles and cytokine release without the confounding effects of spatial variability. Imaging analysis could be expanded through additional replicates to enable co-staining for factors such as IL-8 and TGF-β1 for comparison with LEGENDplex^TM^ results. Further, the LEGENDplex^TM^ analysis of secreted cytokines and growth factors was limited by the high media volume in perfusion systems reducing analyte detectability and quantification accuracy. This analysis could thus be improved by reducing media volume under perfusion conditions and evaluating static cultures under the same dilutions.

The PDMS IA models developed in this study were shown to have no distension under pressure due to the thick-walled block-like structure (Figure S2). Similarly, the compliance of PDMS vascular phantoms have been reported to be lower than numerical simulations and *in vivo* arterial measurements.^79^ While the lack of distension in this study was useful in isolating effects of flow alone, future work should incorporate variable wall thickness models to analyse the impact of biomechanics in combination with WSS. While WSS has been heavily investigated through *in vivo*, *in vitro* and *in silico* studies on IAs, the role of wall biomechanics and remodelling in IA progression remains poorly understood.^23,80,81^ By incorporating wall thickness variability in line with intraoperative observations of the aneurysm wall, the impacts of circumferential strain under intravascular pressure and flow could be investigated.^82^ These studies could be combined with CFD and fluid-structure interaction simulations to evaluate the combined effects of wall thickness, geometry, strain, fluid flow and cellular remodelling on aneurysm rupture prediction.^83^ However, fabrication of these models with a controlled wall thickness can be challenging, with de Nys et al. (2026) describing fabrication methods previously investigated.^84^

## 4 Conclusion

In conclusion, this study demonstrates that endothelialised *in vitro* IA models provide a viable platform for investigating endothelial-haemodynamic interactions relevant to IA growth and stability. Prolonged perfusion beyond 48 hours was required to move beyond acute inflammatory responses associated with flow onset, highlighting an important limitation of existing short-term *in vitro* studies. In patient-specific IA models, regions of high shear impact were associated with reduced endothelial junctional integrity, alongside differential TGF-β1 secretion, IL-8 secretion and FGF-b consumption under cardiovascular flow compared to steady flow. These findings support the utility of endothelial cytokine profiling to monitor endothelial status and remodelling activity in IAs, with potential to complement clinical rupture-risk prediction frameworks. Overall, this work presents the first *in vitro* IA model system that integrates a physiological cardiac flow waveform and captures the temporal endothelial responses over multi-day perfusion.

## 5 Methods

### 5.1 Ethics and patient consent

hTERT immortalized green fluorescent protein (GFP)-expressing human aortic endothelial cells (HAECs) were cultured in accordance with the University of Queensland’s Human Research Ethics committee (2021/HE002698) and the Institutional Biosafety Subcommittee (IBC/602E/ChemEng/2023).

Patient IA images were collected from clinicians at the Royal Brisbane and Women’s Hospital (RBWH) and then fully de-identified before the design and fabrication of IA cell culture models, following RBWH Human Research Ethics Committee (HREC ref: LNR/2019/QRBW/49363), the University of Queensland HREC (ref: 2022/HE001933), and the Queensland Department of Public Health (Public Health Agreement #49363) ^85^.

### 5.2 Segmentation and alignment of intracranial aneurysm with intraoperative images

The IA chosen for fabrication was segmented and modelled as described by Anbananthan et al. (2025) and depicted in **Figure 8** ^86^. Patient Digital Subtraction Angiography (DSA) scans were segmented in Amira (FEI, Hillsboro, Oregon, USA) using pixel intensity thresholding. The inlet and outlet vessels of the model were extended by 15 mm to ensure fully developed flow and to prevent recirculation using ANSYS SpaceClaim (ANSYS Inc, Canonsburg, Pennsylvania) ^86^.

**Figure 8.**
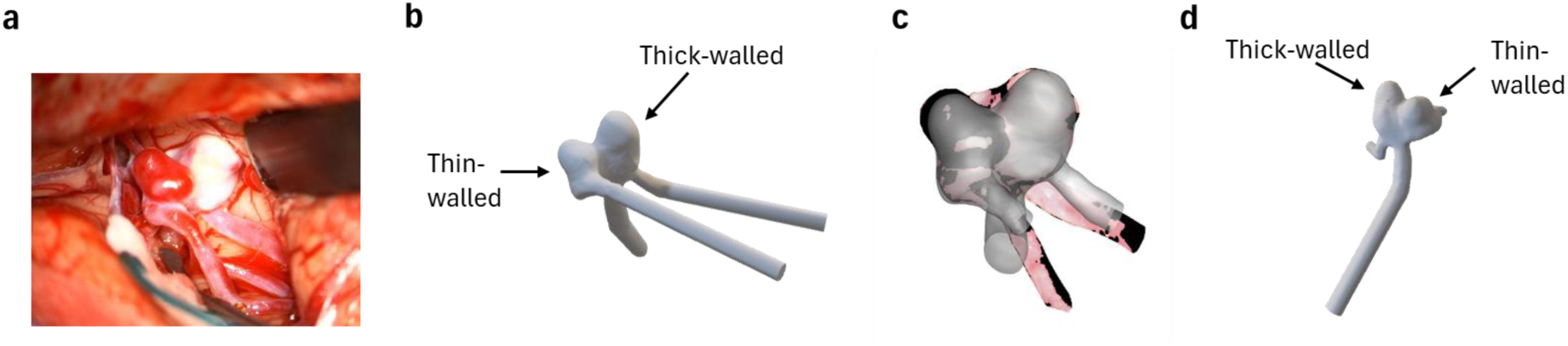
**Intracranial aneurysm morphology and segmentation**. (a) Intraoperative photo of the intracranial aneurysm selected for modelling. (b) Segmented aneurysm with added extensions for flow stabilisation and identification of thin- and thick-walled regions. (c) Alignment of segmented aneurysm against the intraoperative image. (d) Rotation of segmented aneurysm with identification of thin- and thick-walled regions.

### 5.3 Straight channel and intracranial aneurysm model fabrication

3-part moulds for creating idealised channel geometries were created using 3-matic software (Materialise, v25.0), with a central diameter of 2.2 mm, length of 55 mm and an inlet and outlet diameter of 4.1 mm to fit barbed connectors (**Figure 9**a). Moulds were 3D printed in polylactic acid (PLA) using the Ultimaker S3 Fused Deposition Modelling printer, coated with XTC-3D^TM^ print coating (Smooth-On, Inc.) and clamped together.

**Figure 9:**
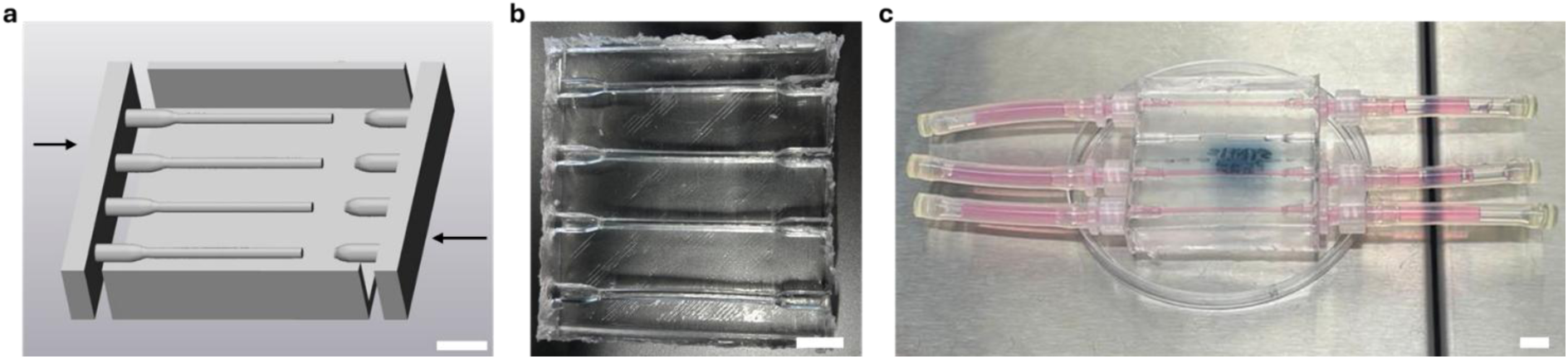
**Idealised PDMS channel design and fabrication**. (a) Computer-aided design (CAD) of 3-part mould, (b) moulded PDMS channels, (c) cellularised PDMS channels connected to 5 cm tubing either side. Scale bars represent 10 mm.

SylgardTM 184 Silicone Elastomer kit (Polydimethylsiloxane or PDMS, Cat. No: 500GM184KIT) was used to prepare PDMS by mixing the Elastomer base with the curing agent in a 10:1 ratio. The mixture was poured into the 3-part moulds and cured at room temperature for 48 hours due to the low melting temperature of PLA, before demoulding (Figure 9b).

PDMS IA models were fabricated using a method previously described ^84^. In brief, an Asiga PRO4K 80 printer (25 µm resolution, 2.4 mW/cm^2^ light intensity) was used to print the sacrificial aneurysm core in a water-soluble (WS) resin (3Dresyns, IM-HDT-WS, Cat.: P20099). Prints were cleaned using the Cleaning Fluid WS1 Bio (3Dresyns, Cat.: P20762) and post-cured using a FormCure (FormLabs) post curing station at 405 nm for 10 minutes at room temperature, submerged in fresh cleaning fluid. External supports were manually removed and models lightly sanded where supports were removed. The water-soluble aneurysm core was then placed into a negative mould and surrounded with PDMS and cured at 80°C for one hour. Once cured, PDMS models were carefully removed from the T-moulds and the sacrificial core dissolved by submersion in water.

### 5.4 Microcomputed tomography (µCT) analysis of fabricated vascular models

PDMS channels connected to barbed luer locks and tubing (1.6 mm inner diameter) were imaged using the Yxlon FF35 CT system to enable computational analysis of fluid dynamics through the channels. The PDMS channel and connectors were exposed to a direct transmission beam at 79.31 kV, 279.88 µA and 22.36 W with an exposure time of 142.86 ms, producing a voxel pitch of 41.67 µm.

Similarly, the fabricated IA models were imaged to evaluate geometric accuracy and distention behaviour as previously described ^84^. Models were exposed to a direct transmission beam at 78.87 kV, 281.51 µA and 22.11 W with an exposure time of 181.82 ms, producing a voxel pitch of 33.33 µm. Scans took approximately eight minutes to complete. The distension behaviour was evaluated by pressurisation with air to 180 mmHg before imaging as previously described ^84^.

Radiographic data was processed using VGSTUDIO MAX (Volume Graphics, v2024.2) and exported in STL format for computational analysis of fluid dynamics and geometric analysis in 3-matic software. To extract the internal geometry, datum plane interactive cuts of the STLs allowed the inner lumen surface of each model to be extracted as a separate part. The resulting models were then aligned with the originally segmented artery to evaluate geometric accuracy, or with the un-pressurized model for analysis of distension. Part comparison analysis was conducted to produce a colour map of geometric differences. The Dice-Sørensen coefficient was then calculated using the Segment Comparison module in 3D Slicer (v5.6.2) ^87^.

### 5.5 Maintenance culture of HAECs

HAECs were routinely cultured in EGM^TM^-2 medium without Gentamicin sulfate-Amphotericin (GA-1000). GA-1000 was added to media when used in 3D material cultures. Cells were thawed at passage 4 and seeded at passage 7 across all experimental conditions and replicates. Cells were cultured at 37°C in a humidified atmosphere of 5% CO_2_ in air.

### 5.6 Seeding HAECs into vascular models

To prepare for cell seeding, vascular models were rinsed with 100% (w/v) isopropanol then distilled water and allowed to dry. Oxygen plasma treatment (70 W, 3 min) was employed to enhance surface activation in these large hollow models, where extended exposure has been shown to improve hydrophilicity.^88^ Channels were then filled with a 2% (v/v) 3-aminopropyltriethoxysilane (APTES, Cat. No: 440140-100ML, Merck Life Science) solution in 100% ethanol for 30 minutes, washed three times with 100% ethanol, lightly dried with compressed air and baked at 100°C for 30 minutes. Samples were then sterilised by soaking in 70% (w/v) ethanol for 10 minutes and washed twice with sterile phosphate buffered saline (PBS) for 5 mins. Channels were filled with 20 µg/mL fibronectin solution (Cat. No: 10838039001, Roche, diluted in DPBS) and incubated at 37°C overnight.

The next day, the fibronectin solution was aspirated from vascular models before cell seeding. With the straight channel models, cells were seeded at 120,000 cells/cm^2^ and then rotated 90° after 30 minutes. After another 30 minutes (60 minutes total), channels were rotated 90° and HAECs re-seeded at the same seeding density. Models were again rotated 90° after 30 minutes. After another 30 minutes (60 minutes total after second seed) media was aspirated to remove unadhered cells and filled with fresh media.

Similarly, HAECs were seeded into the PDMS IA models at 120,000 cells/cm^2^ and models were rotated every 20 minutes onto each of the six flat faces to facilitate even seeding. After the first 60 minutes of seeding on three orientations, HAECs were re-seeded and again rotated every 20 minutes for a total of 60 minutes. Media was then aspirated to remove unadhered cells and filled with fresh media.

Endothelialised models were cultured under static conditions for five hours at 37°C in a humidified atmosphere of 5% CO_2_ in air.

### 5.7 Start-up and continuous operation of perfusion cultures

After five hours of static culture, the cellularised vascular models were connected to a closed, recirculating perfusion circuit comprising a peristaltic pump (Masterflex® Ismatec® Reglo ICC), an empty 60 mL syringe to damp pressure fluctuations and a reservoir with 30 mL media. Media was perfused continuously overnight at 0.2 mL/min to precondition cells. Static cultures in straight channels were connected to 5 cm of extra tubing on either side and 1 mL of fresh media added (Figure 9c). Static PDMS IA models were connected to approximately 5 cm of extra tubing at the inlet of the model for 1 mL total media volume. The next day, flow was automatically ramped linearly between 0.2 mL/min and 31 mL/min over eight hours.

The next day, vascular models were disconnected from the perfusion circuit and 10 mL media was removed for LEGENDplex^TM^ immunoassay and replaced with 20 mL of fresh media. Models were then reconnected to either a peristaltic pump circuit for steady flow or the custom pneumatic cardiac-flow pump as described below and depicted in **Figure 10** and **Figure 11** respectively. All perfusion experiments were performed in triplicate with independent reservoirs.

**Figure 10:**
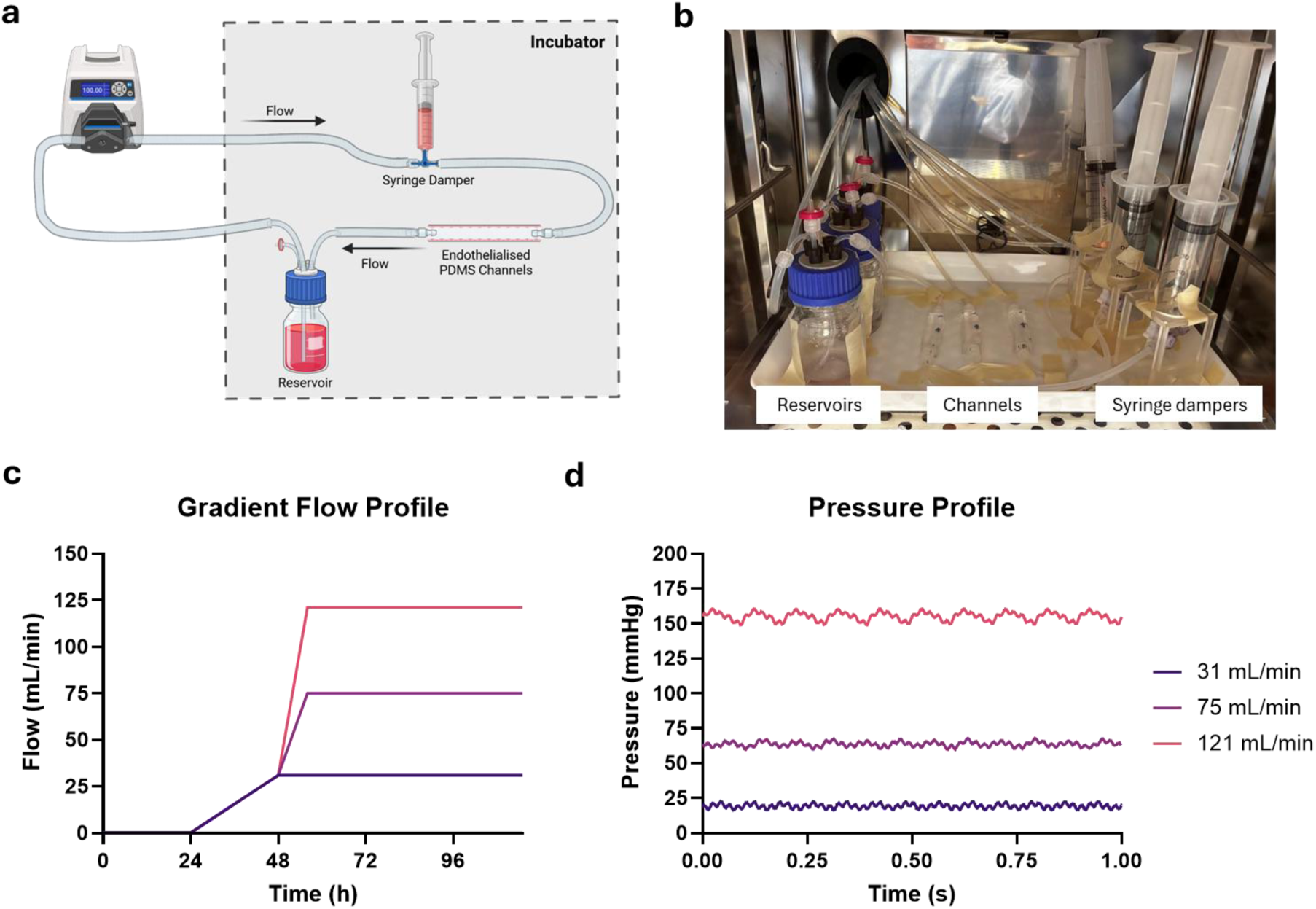
**Straight channel perfusion experimental setup**. (a) Diagram of flow loop with pump external to incubator. Created using Biorender.com. (b) Picture of flow loop inside the incubator. (c) Flow profile. (c) Pressure profile.

**Figure 11:**
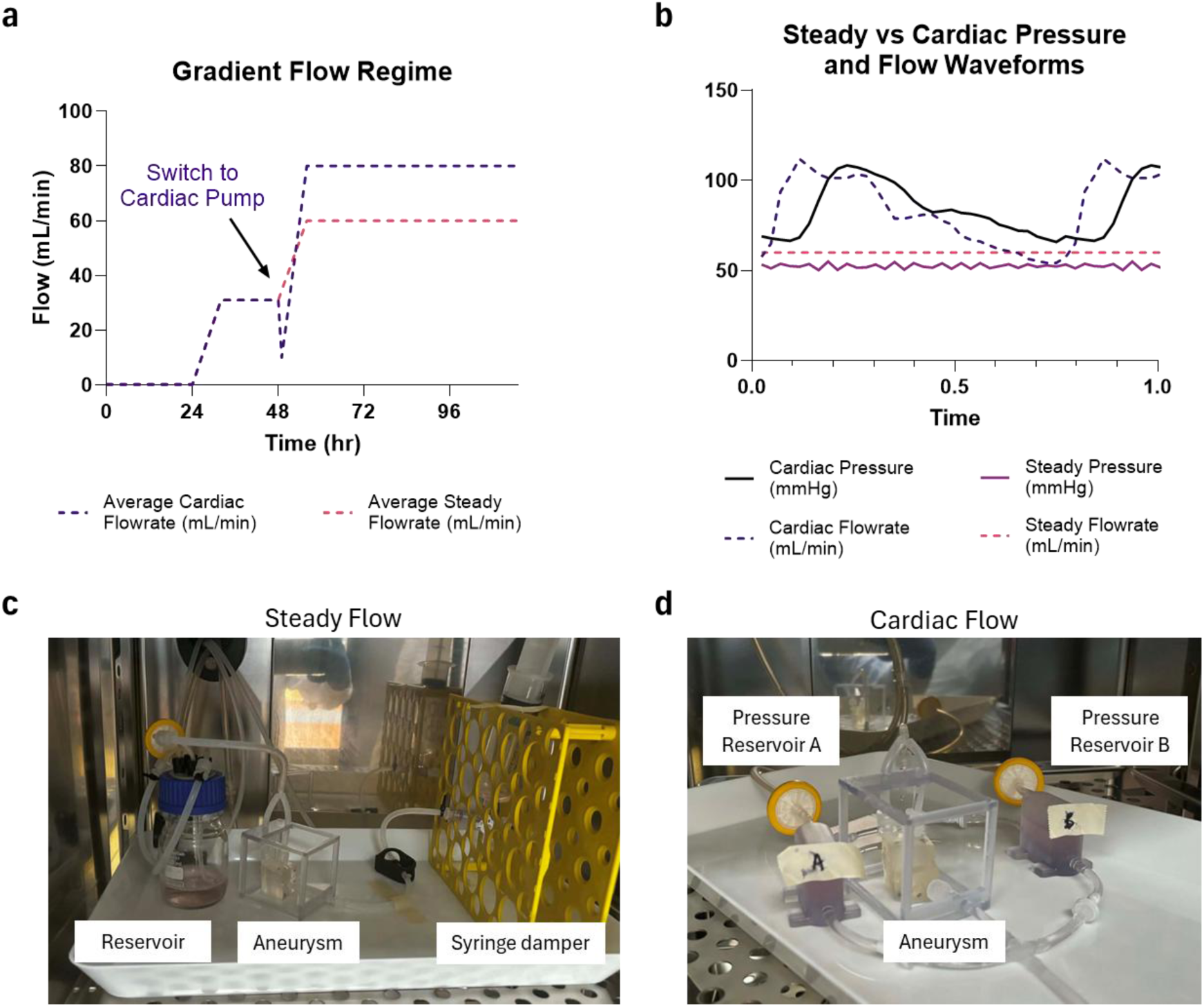
**IA model perfusion experimental setup for steady and cardiovascular flow conditions**. (a) Gradient increases and average flow rate for each flow circuit. (b) Flow and Pressure waveforms. (c) Set-up of steady flow circuit within incubator. (d) Set-up of cardiovascular pump circuit within incubator.

#### 5.7.1 Steady flow perfusion of straight channels

Perfusion was either continued at 31 mL/min for another 48 hrs using the Ismatec® Reglo ICC pump, or tubing was switched to accommodate a larger flow peristaltic pump (Masterflex® L/S®, 07528-10) and ramped further to 75 or 121 mL/min over eight hours. The perfusion system, the gradient flow program and pressure profile is presented in Figure 10. An empty 60 mL syringe was placed upstream as a damping chamber using to reduce pressure pulsations (Figure S1).

At day 5 of perfusion, PDMS channels were disconnected from the fluidic system under aseptic conditions and the first quarter (Q1) of a static and perfused channel cut and stained with Live-or-Dye^TM^ 568/583 viability stain diluted in media (1:1000, Cat. No. 32005, Gene Target Solutions) for 30 minutes at room temperature. All cellularised channels were washed with DPBS before fixation and immunofluorescent (IF) staining. 30 mL of media from each reservoir was sampled for immunoassay.

To demonstrate the retention of the fibronectin coating under perfusion, PDMS channels were coated with fibronectin and exposed to 121 mL/min flow of PBS in the absence of cells. Following perfusion, channels were incubated with rhodamine-B isothiocyanate (RBITC, 20 µg/mL, Cat. No.: R1755, Sigma-Aldrich) as a fluorescent stain to visualise surface-associated protein. Given the absence of cells, the observed signal is expected to predominantly reflect retention of the fibronectin coating. Results are presented in Figure S2.

#### 5.7.2 Perfusion of IA models

For the steady flow regime, flow was automatically ramped at day 2 linearly between 31 mL/min to 60 mL/min over 8 hours. Flow was maintained at 60 mL/min until day five (Figure 11).

A custom pneumatic cardiovascular pump was used for generating a cardiac flow regime ^89^. A physiological flow waveform of the right middle cerebral artery (MCA) velocity was used as the basis for this system, representing patient ID s0030 as published by Novak et al. (2009) ^90,91^. Flowrate (mL/min) was calculated based on an inlet diameter of 2.2 mm of the chosen MCA aneurysm model. Average flowrate was 146 mL/min, consistent with literature reports of the flowrate through the MCA ^92^. Flow was scaled down to the average flowrate through distal branches of the MCA of 43 mL/min ^92^. This waveform was used for blood-flow CFD simulations. Because culture media is less dense and less viscous than blood, flowrate was scaled up to an average of 80 mL/min to achieve a similar experimental WSS profile (Figure 2).

At day two, the aneurysm models were disconnected from the perfusion circuit, 10 mL sampled for LEGENDplex^TM^ and supplemented with 20 mL fresh media. IA models were then connected to the cardiovascular pump perfusion circuit and filled with 16 mL of the media mixture. When transitioning to the cardiovascular pump, an average flow of 10 mL/min and an average pressure of 30 mmHg were first applied to mitigate cell shock that could be induced by the introduction of pulsatile flow. This was manually increased every hour for eight hours until an average flow of 80 mL/min and 80 mmHg was achieved. Flow was maintained at this rate until day five of perfusion. The frequency of oscillations was set at 80 beats per minute. The perfusion system, the gradient flow program and pressure profile is presented in Figure 11.

### 5.8 Computational Fluid Dynamics (CFD) analysis

CFD simulations were performed in ANSYS Fluent (v2024.R1) assuming incompressible, laminar flow. For the straight channel simulations, three flow rates (31, 75, and 121 mL/min) were simulated on a locally refined mesh (minimum/maximum local cell length of 0.05 mm to 0.2 mm to capture diameter changes) with 8 inflation layers. Because residual reduction was more challenging at the highest flow regime, solution acceptance combined residual monitoring with a mass-conservation check. Momentum residuals were on the order of 10^-5^ and continuity residuals 10^-2^, while the inlet–outlet mass-flow imbalance remained within 0.15% of the imposed inlet flow (−0.071%, −0.00529%, and 0.144% for 31, 75, and 121 mL/min, respectively).

For the IA models, flow was simulated on a locally refined mesh where the domain was limited to an element size of 0.1 mm and 8 inflation layers at the walls to resolve near-wall gradients. The simulation was run for two cardiac cycles using 1032-time steps in total. Convergence within each time step was assessed using scaled residual targets of 10^-5^ for continuity and 10^-6^ for momentum. Simulations using culture media were modelled with a fluid density of 1002 kg/m^3^ and dynamic viscosity of 0.0007 kg/m^3^ ^93^. Simulations of blood flow were modelled with a fluid density of 1060 kg/m^3^ and a dynamic viscosity of 0.004 kg/m^3^ ^94^. Wall shear stress (WSS) and velocity contours were created and exported.

### 5.9 Immunofluorescent staining

Samples were fixed in 4% paraformaldehyde (Cat. No: C006, ProSciTech Australia). Samples were then washed three times with PBS, permeabilised with 0.1% (v/v) Triton^TM^ X-100 solution (Cat. No: 93443, Sigma-Aldrich) for 10 minutes, washed with PBS and then blocked using 10% (v/v) goat serum (Cat. No: 16210064, Thermo Fisher Scientific) in PBS for 60 minutes at room temperature. Channels were sliced in half lengthwise, with one half to be stained with primary antibodies and the other with the corresponding isotype. Large, stitched fluorescence images were acquired of Q2-4 using a 4x objective on a Nikon Eclipse Ti2 fluorescence microscope and the ‘Scan Large Image’ module in the NIS-Elements software with automated z-series capture. Channels were then cut into four sequential quarters (Q1-4) along their length to enable staining of different IF panels across the channel. IA models were sectioned such that each segment of the asymmetric aneurysm geometry could be sliced in half, with the bottom of each aneurysm segment used for tiled-fluorescent imaging as performed on the straight channels.

IF staining was performed in the dark using primary antibodies and isotypes detailed in Table S1 for three hours at room temperature. Samples were washed three times with PBS, then stained with secondary antibodies as detailed in Figure S3 for one hour. All samples were stained with NucBlue^TM^ Fixed Cell ReadyProbes^TM^ Reagent (2 drops per mL in DPBS, Hoechst 33342, Cat. No: R37605) at room temperature for 30 minutes. The Q4 straight channel segment was also stained with ActinRed^TM^ 555 ReadyProbesTM Reagent (2 drops per mL in DPBS, Cat. No: R37112) at room temperature for 60 minutes. Similarly, the IA models were stained with ActinGreen^TM^ 488 ReadyProbesTM Reagent (2 drops per mL in DPBS, Cat. No: R37110) at room temperature for 60 minutes.

Fluorescence images were acquired using a Nikon Eclipse Ti2 fluorescent microscope using a 10x objective with z-series automated capture.

### 5.10 Immunofluorescent image analysis

All z-series stacks captured on a 10x objective were initially processed in ImageJ with a background subtraction, the Stack Focuser plugin to create a focused projection, and a linear brightness and contrast adjustment ^95^. All analyses were conducted in Fiji (ImageJ) using custom automated macros to ensure consistent processing across all images ^96^.

Mean fluorescence intensity (MFI) was calculated as the average pixel intensity on samples stained for IL-6, eNOS, CD-31, ZO-1 and Hif-1α, and a pooled isotype MFI for each marker was subtracted to correct for non-specific signal and background fluorescence.

Negative values indicate signal is below background and therefore considered undetectable. As bright cells or spots can cause fluorescent halos in focused projections, these regions of interest (ROIs) were manually selected and excluded from analysis. Using masks created from DAPI and FITC (GFP-fluorescing HAECs) images, nuclear and cytoplasmic MFI respectively were quantified for IL-6, eNOS and Hif-1α expression. Where Q1 was stained for dead cells using the Live-or-Dye^TM^ 568/583 viability stain, viability (%) was calculated using Equation 1 below.

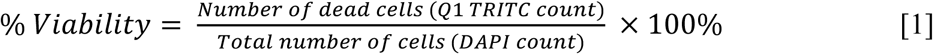

Similarly, the percentage of proliferating cells was determined using Equation 2 below.

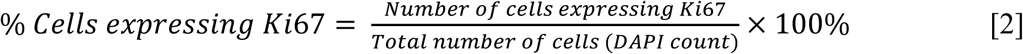

Cell alignment was quantified from CD31-stained images using the OrientationJ plugin ^97^. Alignment strength (R) was calculated based on the Rayleigh test of circular statistics using Equation 3 below, where orientation angles were summarised in a 180-bin histogram (1° resolution), and for each bin *i*, denotes the bin angle and the number of pixels with orientations in that bin ^98–100^. CD31 and ZO-1 colocalisation was calculated using the Fiji (ImageJ) Colocalization Threshold plugin and the Pearson’s correlation coefficient reported ^96,101^.

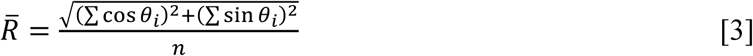

### 5.11 LEGENDplex^TM^ immunoassay

After perfusion, 10 mL of media from each reservoir was centrifuged at 2000 RCF for 10 minutes at room temperature, and 30 µL aliquots of the supernatant frozen at -80°C. The concentration of thirteen cytokines was analysed in the supernatant using the LEGENDplex^TM^ Human Inflammation Panel 1 (13-plex, Cat. No.: 740809, Lot. No.: B452936, BioLegend), including interleukins (IL) IL-1β, IL-6, IL-8, IL-10, IL-12p70, IL-17A, IL-18, IL-23, IL-33, IFN-α2, IFN-γ, TNF-α and MCP-1. The concentration of 13 growth factors was analysed in the supernatant using the LEGENDplex^TM^ Human Growth Factor Panel (13-plex, Cat. No.: 741282, Lot.No.: B467878, BioLegend), quantifying levels of TGF-β1, Angiopoietin-2, EGF, EPO, G-CSF, GM-CSF, HGF, M-CSF, PDGF-AA, PDGF-BB, SCF, FGF-basic and VEGF.

The assay was performed as per the manufacturer’s instructions and analysed using a CytoFLEX S (Beckman Coulter). Instrument configuration was optimized using the LEGENDplex™ Setup Beads, which were run to establish appropriate photomultiplier tube (PMT) voltages for bead classification and PE-based reporter detection. Voltages were adjusted until bead populations were clearly resolved and positioned on scale within the designated plots. Acquisition was monitored in real time to verify stability of bead populations, event rates, and signal distribution. Stopping criteria were based on predefined event counts to ensure sufficient bead representation per analyte. All datasets were saved as raw FCS files and analysed using the LEGENDplex™ Data Analysis Software Suite.

For each panel, samples were analysed across three separate LEGENDplex^TM^ runs with standards. Limits of detection (LOD) and limits of quantitation (LOQ) were therefore reported for each run. Values above LOD but below the LOQ were included in the analysis but considered less precise. Analyte concentrations were corrected for differences in media volume and media replacement between static and perfusion cultures. Media-only controls (1 mL) placed inside the incubator for the duration of the experiments were also sampled at day zero, two and five and used to correct concentrations prior to volume normalisation. Where growth factors were already present in the endothelial cell media, cell-related consumption or secretion rate of growth factors were calculated using Equations 4 to 9, where C is concentration (pg/mL) and D is day.

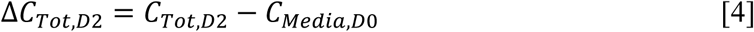

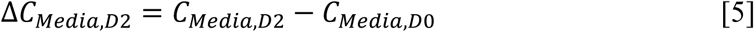

Equation 6 was used to calculate cell-only related changes in growth factor concentration.

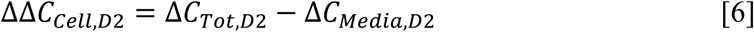

Therefore, the day 2 cell-only concentration (pg/mL) can be calculated using Equation 7.

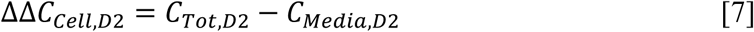

Rate of change (pg/day) was then calculated based on the duration of flow exposure in days and the volume measured at day two. After correction for media removal from sampling and the addition of fresh media, the new mass of each analyte and the resulting concentration was calculated. To calculate the cell-related change in analyte concentration between day two and five, Equation 8 was used.

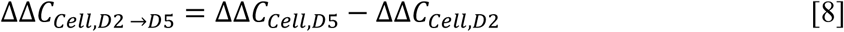

Rate of change (pg/day) was then calculated using Equation 9 to correct for volume differences and duration of flow exposure. These rates were then normalised against average cell density measurements determined at day five (Figure 3) averaged across the model.

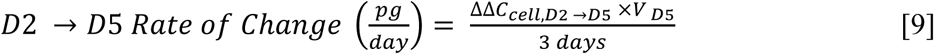

An example of the raw measurements and calculations applied for MCP-1 detected using the LEGENDplex^TM^ Human Inflammation Panel 1 is presented in Supplementary Data 19 and in Supplementary Data 20 for FGF-b detected using the LEGENDplex^TM^ Human Growth Factor Panel.

### 5.12 Statistical Analysis

IF studies were analysed for three biological replicates per condition each with two technical imaging replicates. Similarly, LEGENDplex^TM^ immunoassay was conducted for three biological replicates per condition and analysed in duplicate.

Statistical analysis of cell density measurements was performed using a two-way analysis of variance (ANOVA) with Tukey correction and pairwise comparisons using GraphPad Prism (v10.2.0). Similarly, statistical analysis on immunofluorescent measurements was performed using an unmatched one-way ANOVA with Tukey correction and pairwise comparisons. For LEGENDplex^TM^ results, statistical comparisons were only performed where measurements were above the limit of quantitation (LOQ). Where flow conditions were identical at day two, LEGENDplex^TM^ results for all flow samples were combined at this timepoint and an unpaired two-tailed t-test with Welch’s correction due to the uneven sample size was performed. The same analysis was also performed to compare the analyte rate of change in straight channel experiments under 31 mL/min between day two and five. Day five LEGENDplex^TM^ results were analysed using a one-way ANOVA with Tukey correction and pairwise comparisons. Results are expressed as a mean ± standard deviation, with p<0.05 accepted as statistically significant and indicated using a * (*p<0.05, **p<0.01, ***<0.001, ****<0.0001).

## Acknowledgements

The authors would like to thank the Royal Brisbane and Women’s Hospital (RBWH) Foundation and the RBWH Department of Medical Imaging for their support, including RBWH Foundation grants in 2020 and 2021. The authors would also like to thank Metro North Health who funded materials and some of the researchers through the CranioFacial program of the Herston Biofabrication Institute. This work was supported by funding from the Queensland–Bavaria Collaborative Research Program (QLDBAVDEV20240024), the National Heart Foundation of Australia (108132-2024_FLF, 108420-2024_VG), the Australian Research Council (DE220100757), NSW Health in the form of NSW Senior Research Grant and Advance Queensland (AQIRF1312018). Domestic and international collaborations were supported by an Australasian Society for Biomaterials and Tissue Engineering domestic travel grant and an Emeritus Professor Ted White AM Development Scholarship.

This work also used the Queensland node of the NCRIS-enabled Australian National Fabrication Facility (ANFF) and the microCT available through the Queensland Centre for Advanced Imaging, part of the National Collaborative Research Infrastructure Strategy (NCRIS) and the National Imaging Facility (NIF), with financial contributions from the Commonwealth Government of Australia, the Queensland State Government and The University of Queensland (UQ).

The authors would like to thank: Roozbeh Fakhr, Liam Georgeson, Lauren Drabwell and Dr David Forrestal from the Herston Biofabrication Institute for their assistance in 3D modelling, printing and processing, Professor James Vaughan from UQ for provision of their Masterflex® L/S® peristaltic pump that enabled these high-flow experiments, Malcolm Marker from UQ for engineering a control system with LabView program for this high-flow peristaltic pump to enable automated gradient flow control, Dr Ekaterina Strounina and Associate Professor Gary Cowin from the University of Queensland Centre for Advanced Imaging, for their assistance with µCT imaging and processing, and Professor Justin Cooper-White and Ms Taryn Smith from the University of Queensland for the use of their pressure transducer and for aiding pressure monitoring studies.

We acknowledge the use of M365 Copilot to assist with the polishing of this manuscript to improve readability. Following the use of this tool, all authors reviewed and edited the content for accuracy and intellectual integrity.

## Supplementary Data 1: Pressure damping using upstream syringe

The 60 mL syringes placed upstream reduced peristaltic pump pulsation amplitude by 90% when running at 35 mL/min.

**Figure S1:**
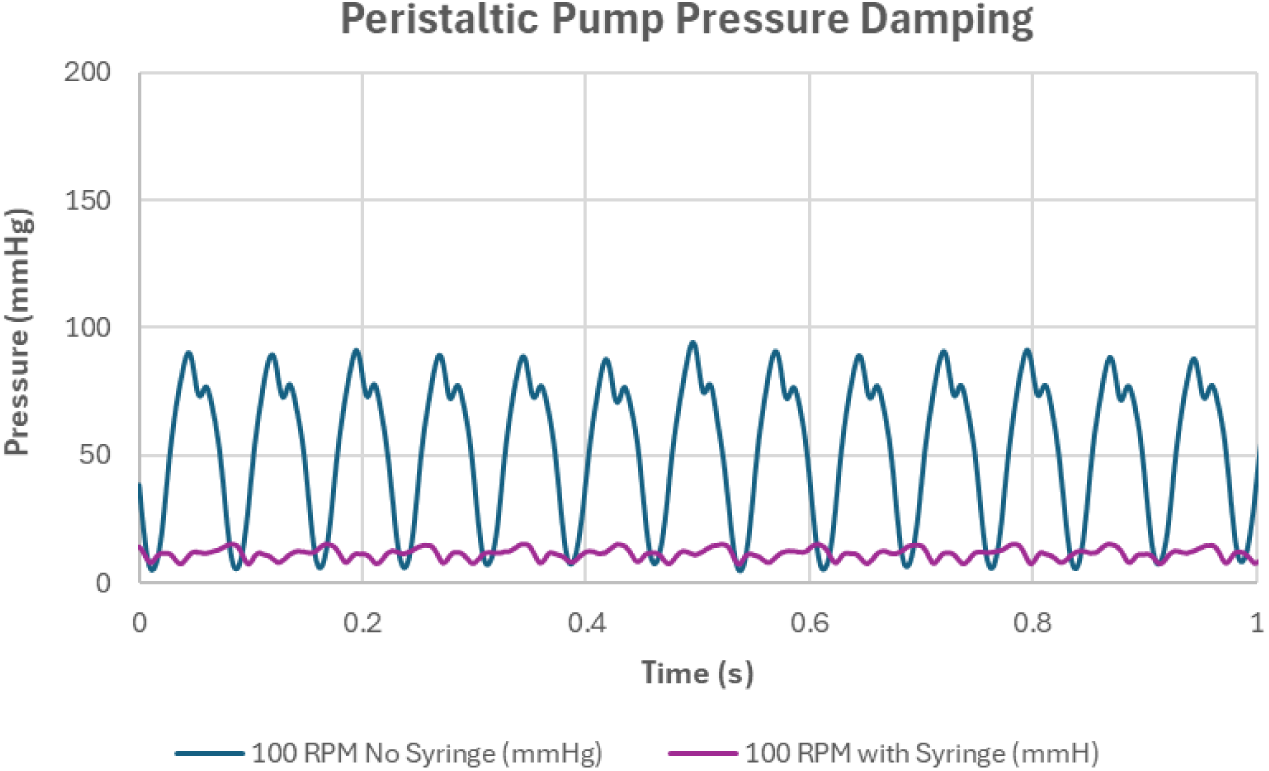
Effect of upstream open 60 mL syringe space buffer on pressure profile from a peristaltic pump.

## Supplementary Data 2: Fibronectin Coating

Visualisation of fibronectin coating on PDMS after perfusion by rhodamine-B isothiocyanate staining (Red).

**Figure S2:**
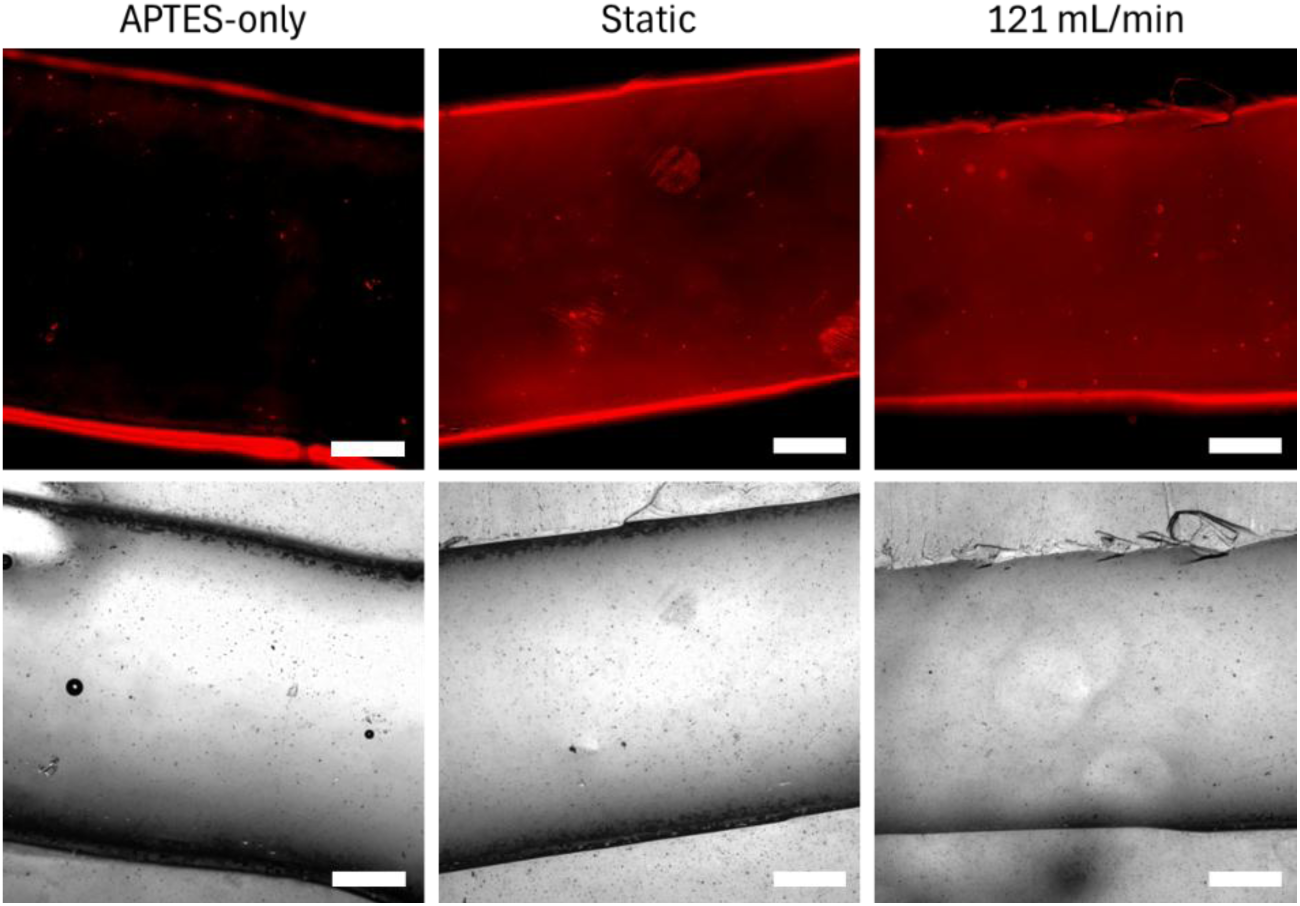
Rhodamine B isothiocyanate (RBITC) staining of fibronectin coated PDMS channels under flow. Scale represents 500 µm.

## Supplementary Data 3: Primary and secondary antibody details

All antibodies were manufactured by Invitrogen and supplied by ThermoFisher Scientific. All secondaries were prepared to a concentration of 4 µg/mL.

**Table S1:**
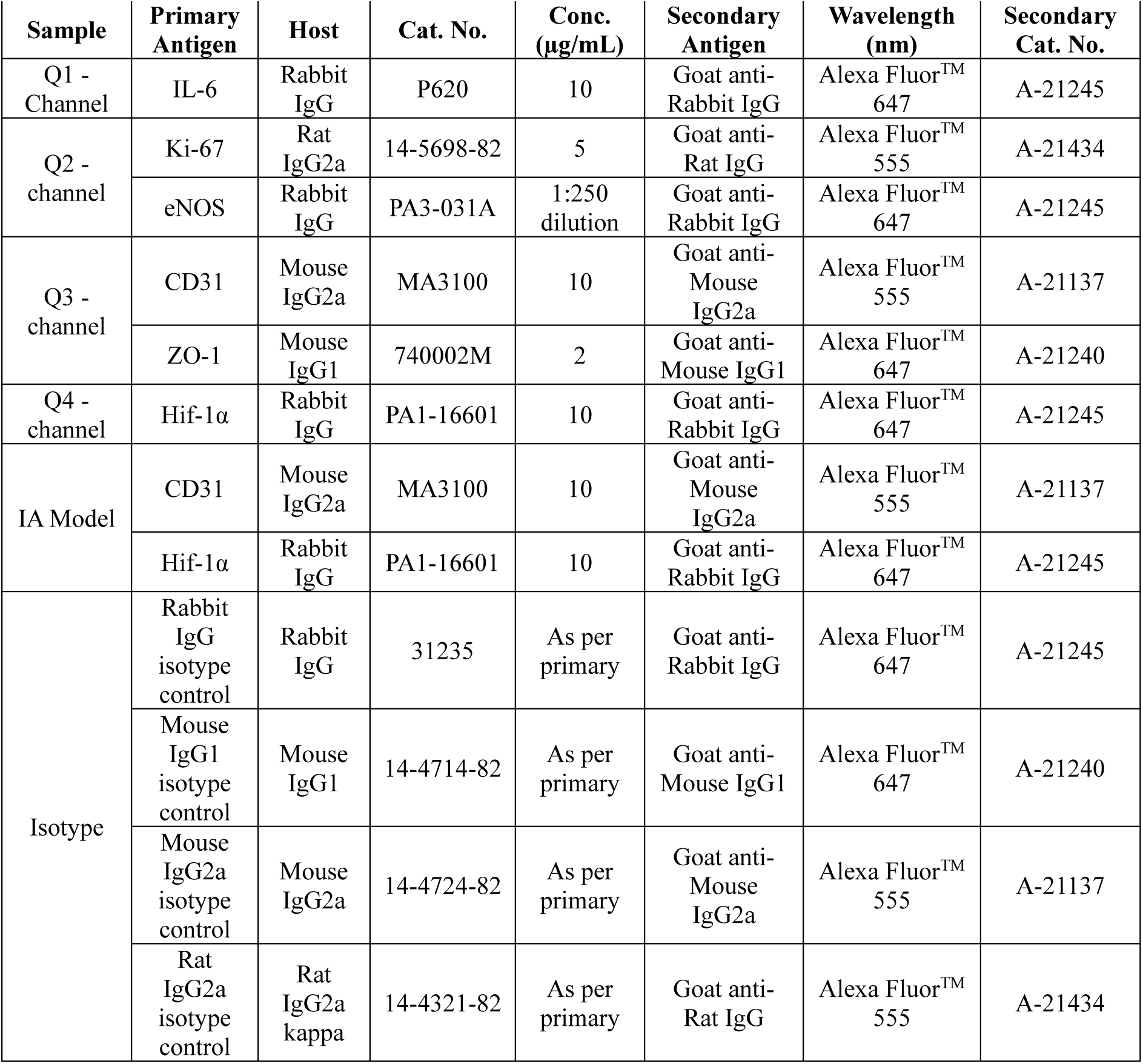
Primary and secondary antibody details.

## Supplementary Data 4: Accuracy and Distension Profile

**Figure S4:**
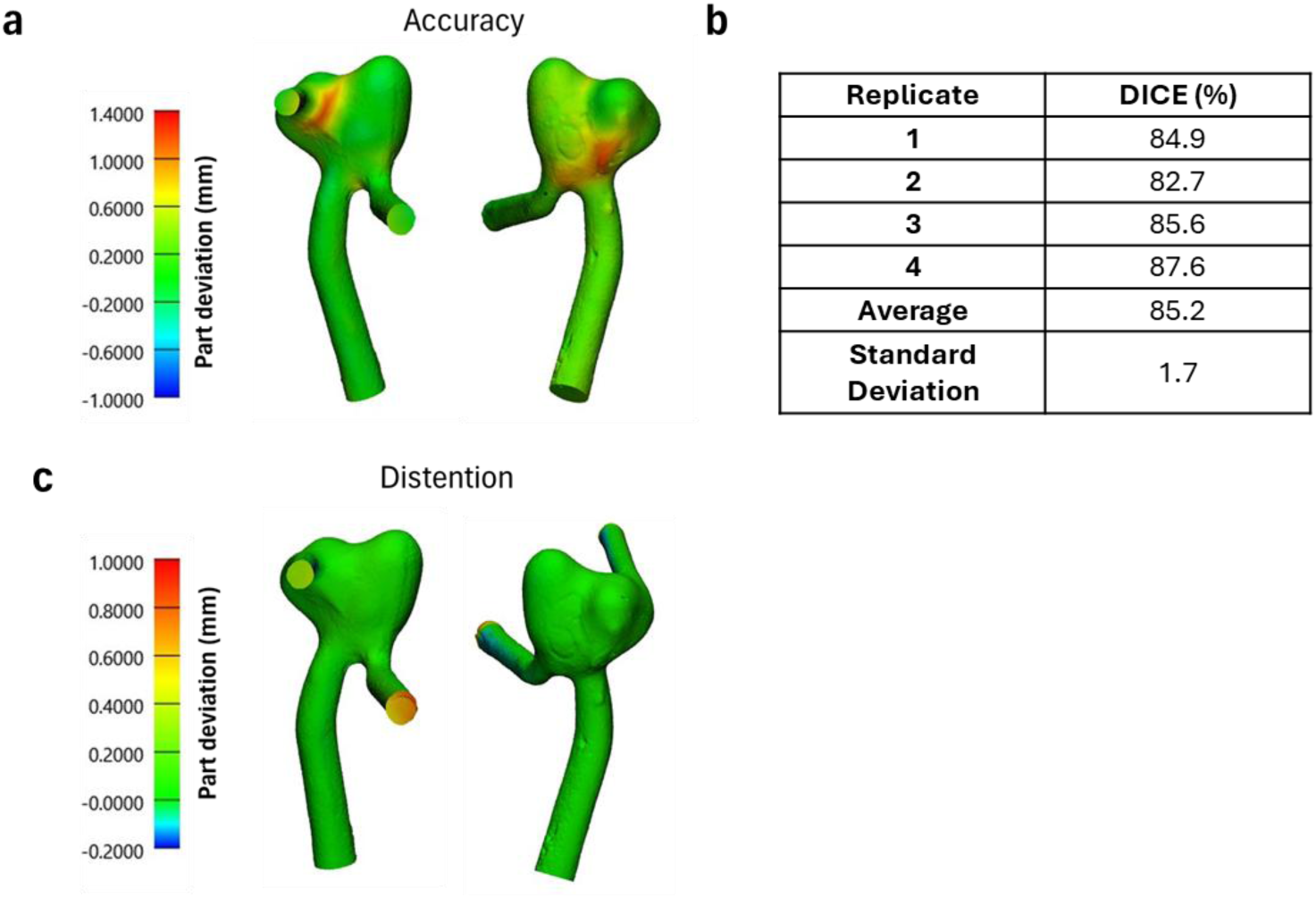
Accuracy and distension analysis of fabricated patient-specific IA models. (a) Colour map of part deviation of one representative IA model. (b) DICE scores for four fabricated IA models. (c) Colour map of part deviation of IA models pressurised to 180 mmHg.

## Supplementary Data 5: Spatial analysis of wall shear stress across aneurysm models

**Figure S5:**
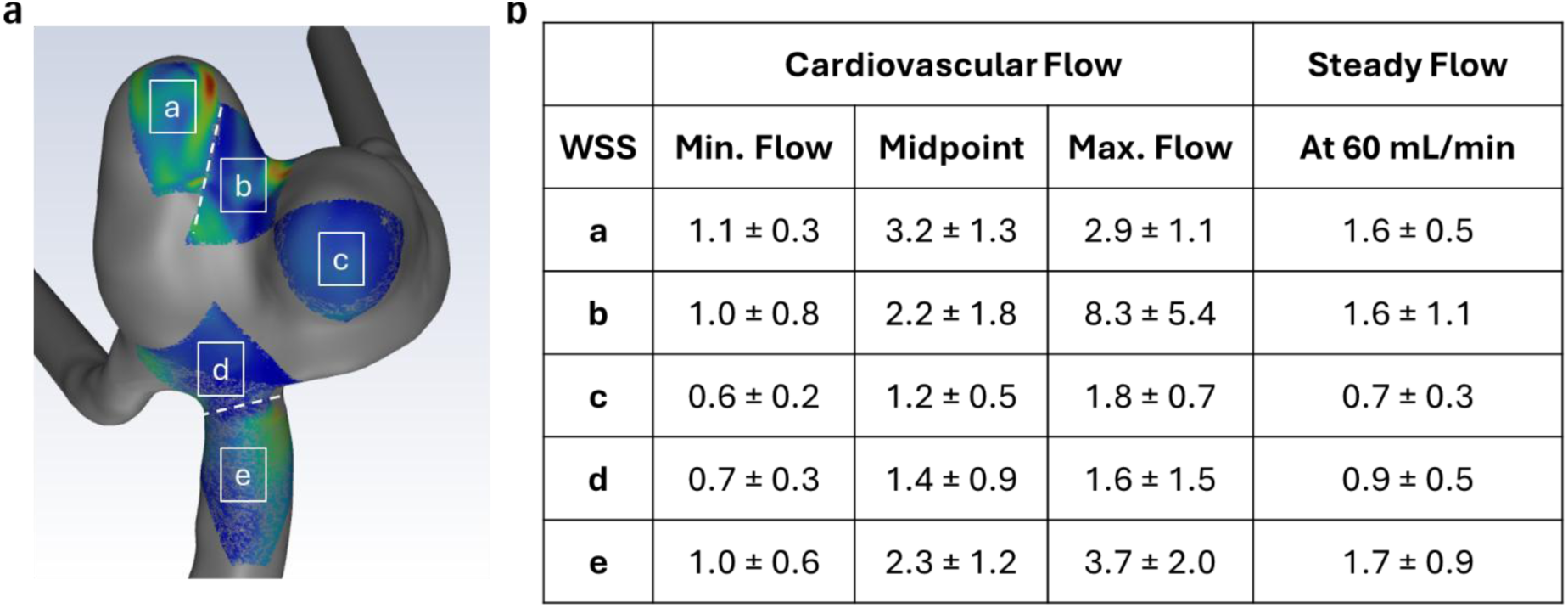
Spatial analysis of wall shear stress (WSS) from the computational fluid dynamics analysis. Locations of analysis labelled ‘a’ to ‘e’ correspond to regions used for subsequent image analysis. (a) Regions of interest (ROI) used for analysis of WSS. (b) Average WSS calculated for each ROI under cardiovascular and steady flow regimes.

## Supplementary Data 6: Q1 Immunofluorescent staining of dead cells and IL-6 on straight channels

**Figure S6:**
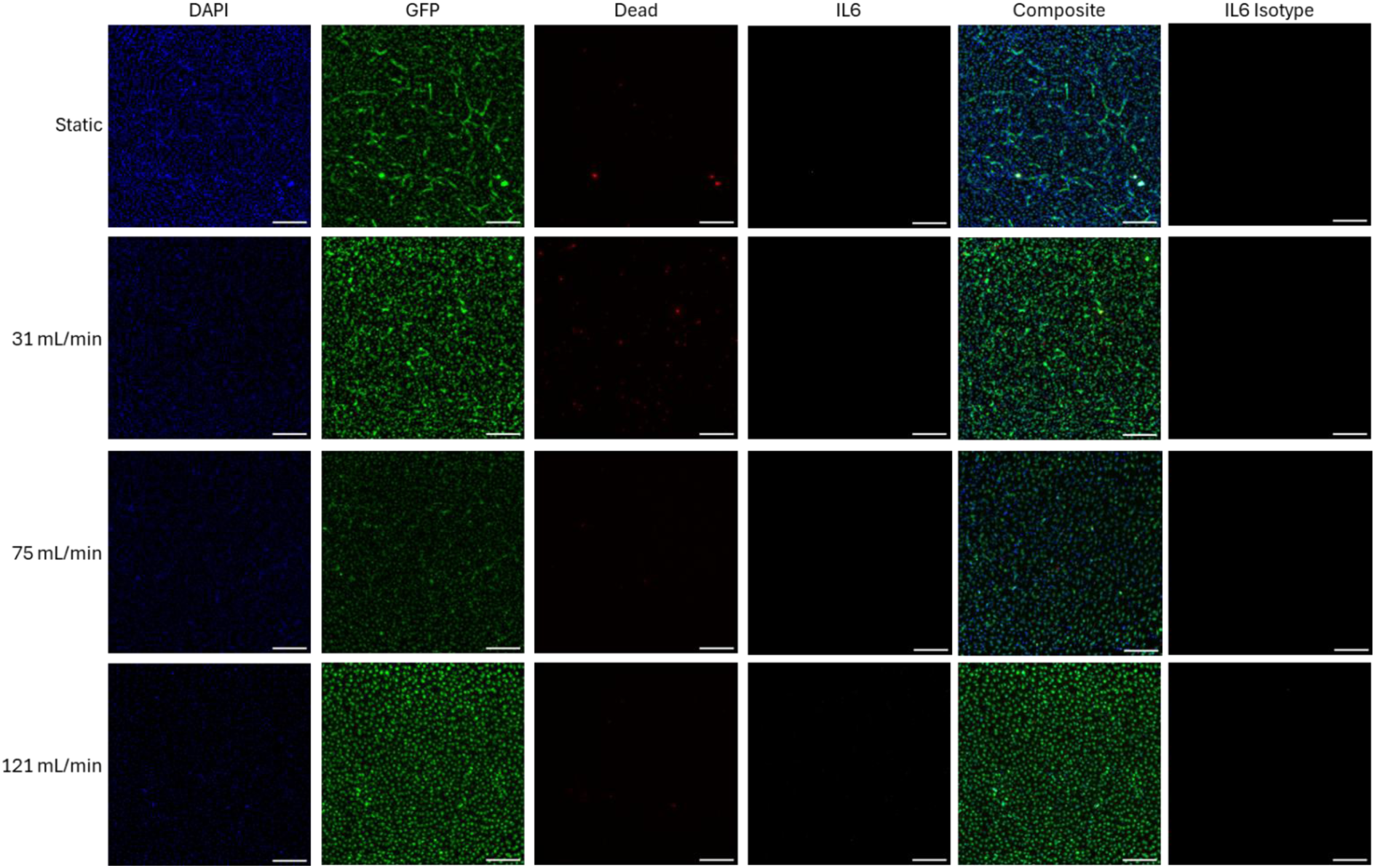
Immunofluorescent staining of human aortic endothelial cells (HAECs) on Q1 of polydimethylsiloxane (PDMS) channels under static and perfused conditions. Stained for DAPI (blue), viability stain (red), anti-IL6 antibody (magenta). Cells express green fluorescent protein (GFP, green). Scale represents 200 µm.

## Supplementary Data 7: Tiled immunofluorescent imaging of HAEC coverage across IA replicates following perfusion

**Figure S7:**
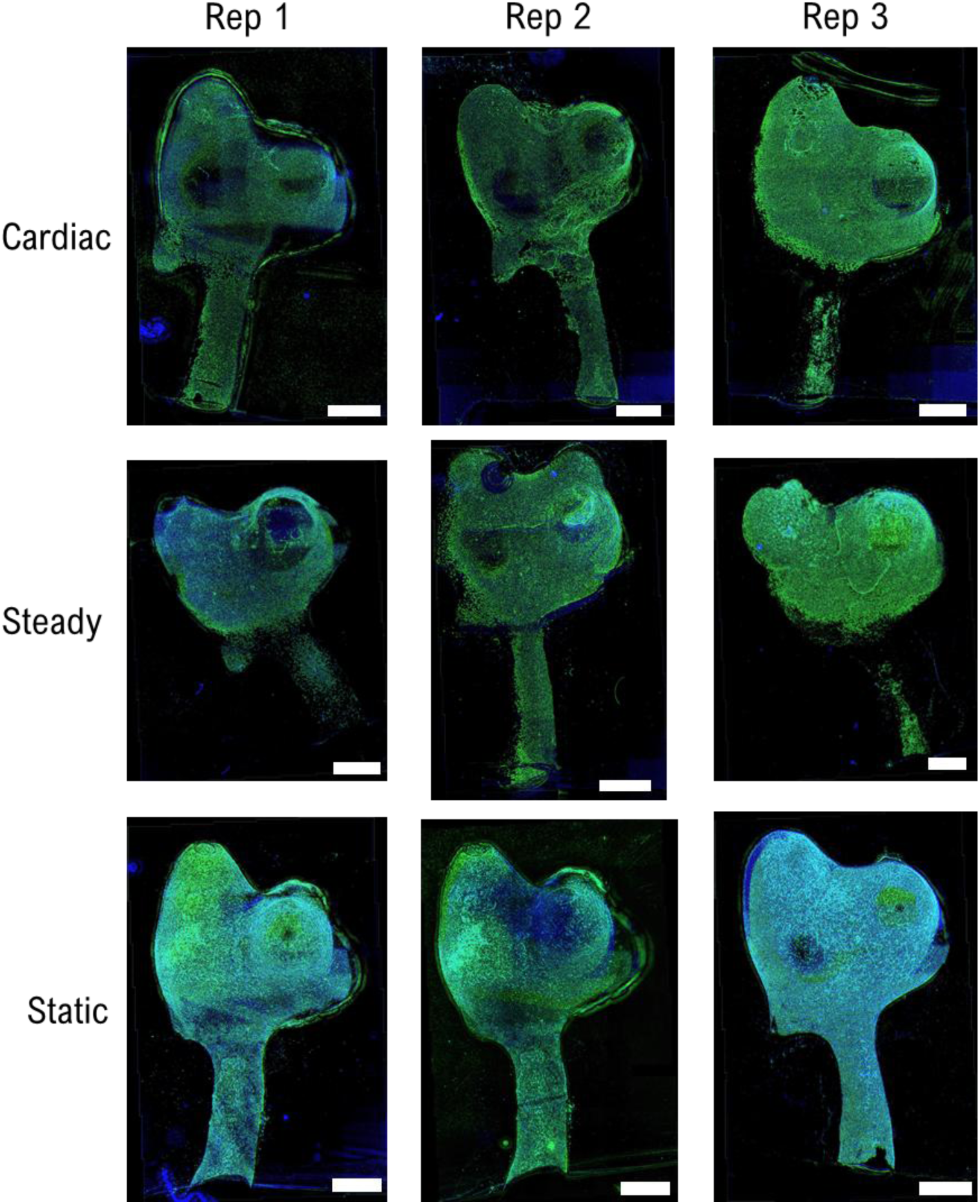
Human aortic endothelial cells (HAEC) on *in vitro* polydimethylsiloxane (PDMS) models of a middle cerebral artery aneurysm after five days of perfusion under cardiac and steady flow profiles. HAECs stained for F-actin (green) and DAPI (blue). Scale represents 2 mm.

## Supplementary Data 8: Effect of wall shear stress magnitude on cell density

Investigating the effect of wall shear stress magnitude on cell density normalised to static conditions. Combines results from straight channel and IA model perfusion studies.

**Figure S8:**
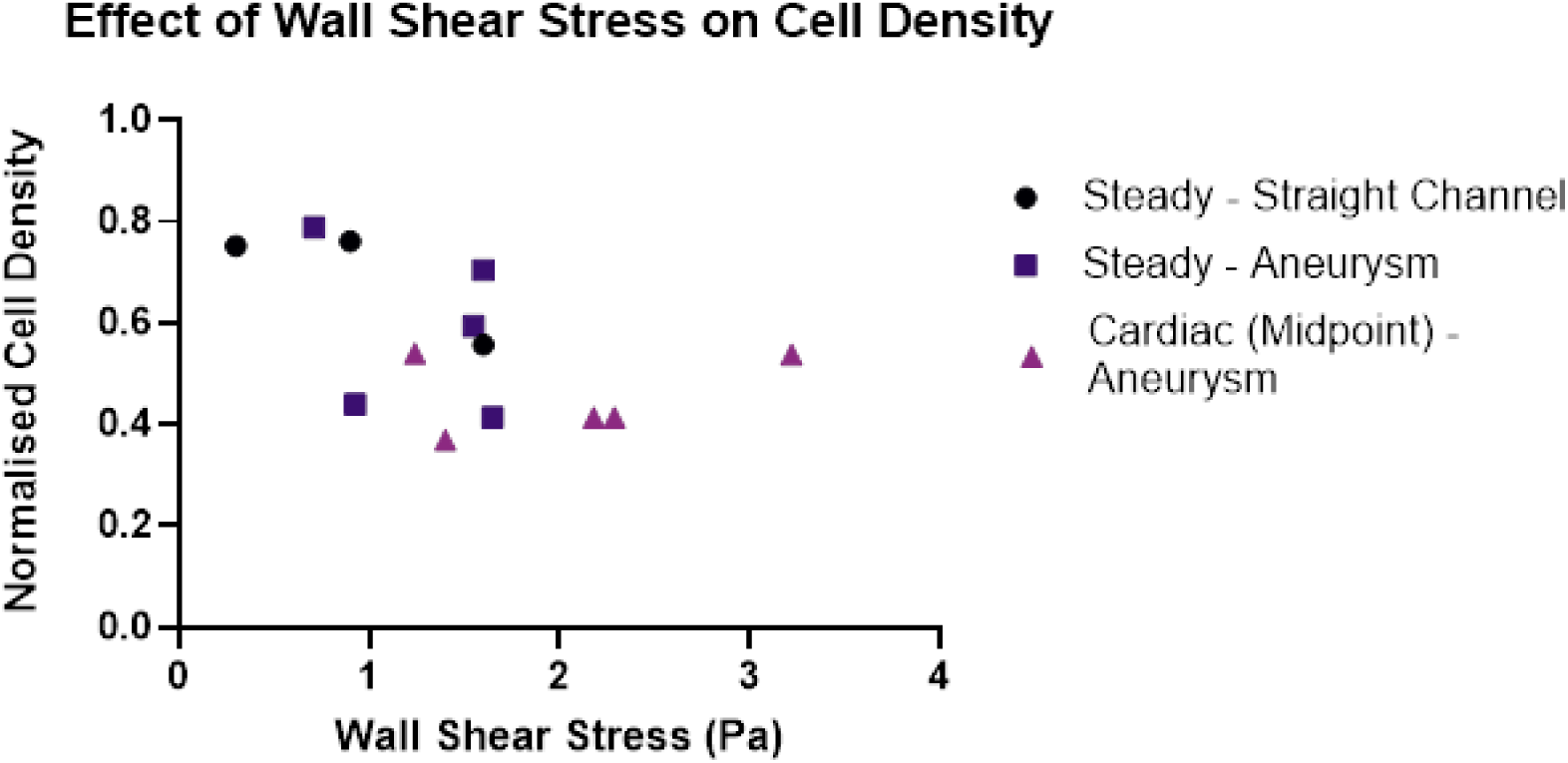
Effect of wall shear stress magnitude on human aortic endothelial cell (HAEC) cell density normalised to static controls in in vitro PDMS models.

## Supplementary Data 9: CD31 and ZO-1 staining with isotypes

**Figure S9:**
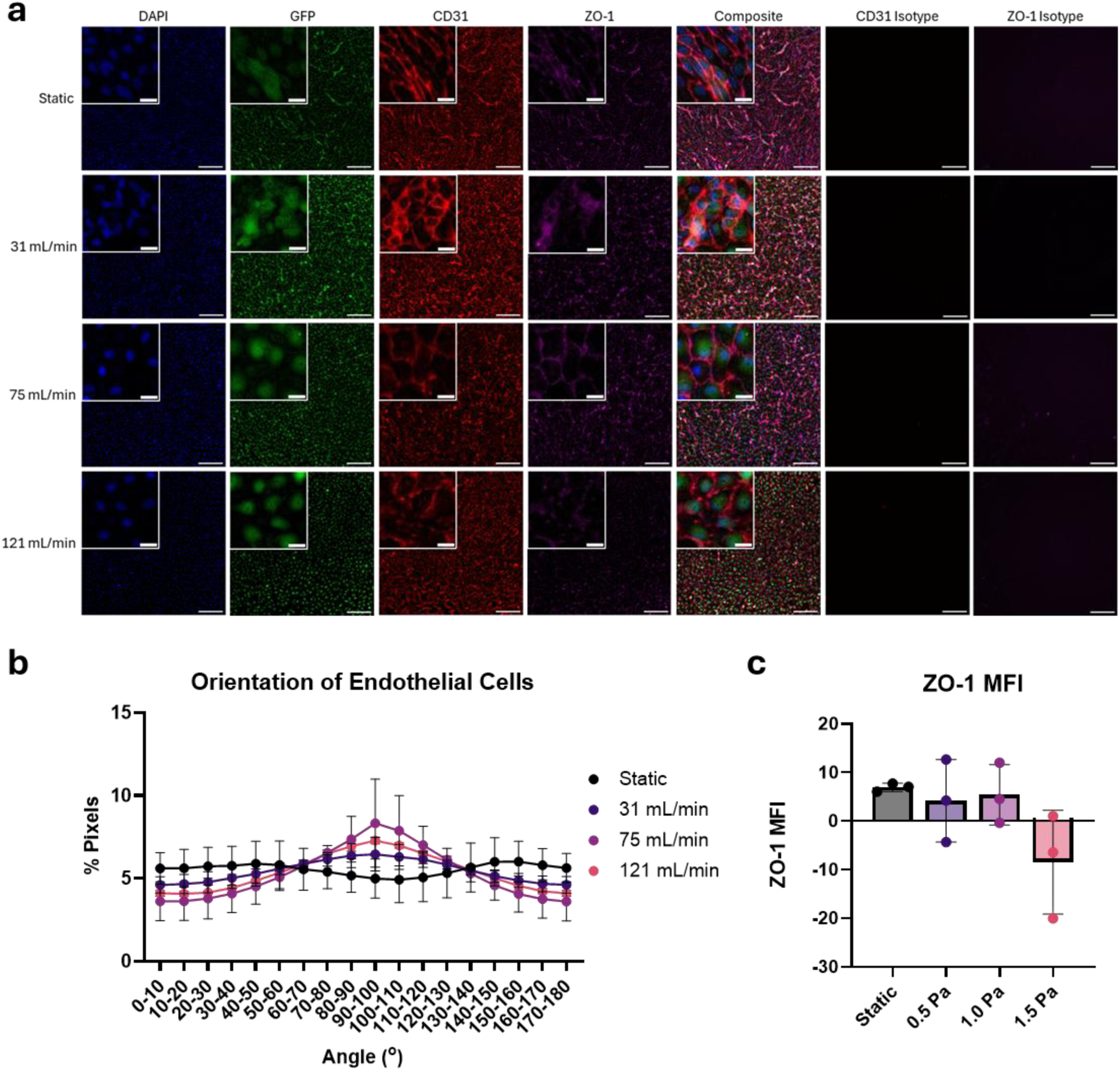
CD31 and ZO-1 expression in human aortic endothelial cells (HAECs) under static and flow conditions. (a) Immunofluorescent images of GFP (green) fluorescing HAECs on PDMS channels stained for DAPI (blue), CD31 (red) and ZO-1 (magenta). Scale of embedded image represents 20 µm. Scale of larger low-resolution image represents 200 µm. (b) CD31 pixel orientation analysis at day five. (c) Mean fluorescence intensity of ZO-1 corrected against isotype controls.

## Supplementary Data 10: Endothelial morphology over time with quantification of orientation angle and ZO-1 MFI at day 5

**Figure S10:**
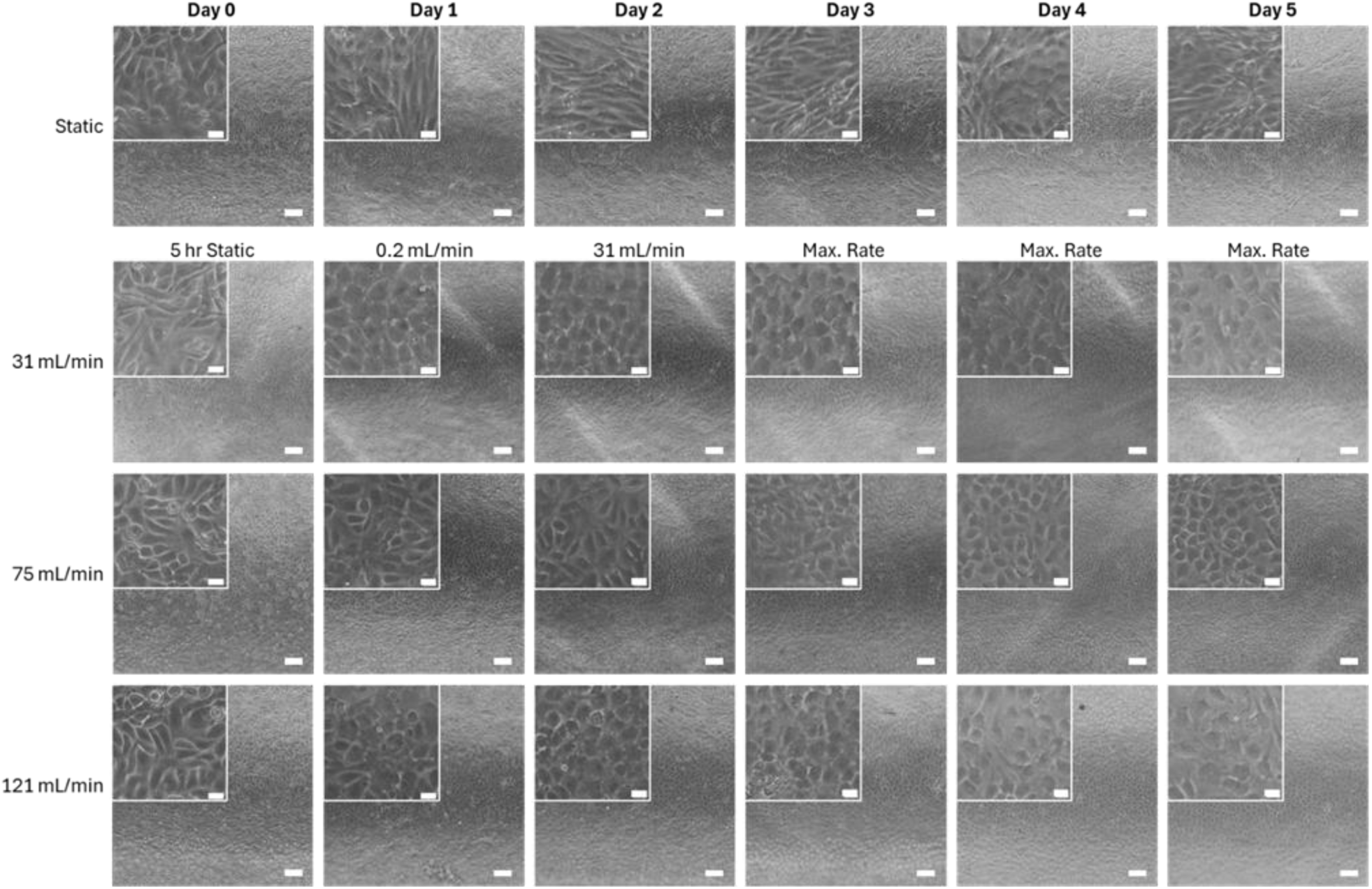
Brightfield images of HAECs over five days of perfusion at 31 mL/min, 75 mL/min and 121 mL/min compared to static conditions. Cells exhibited an elongated morphology under static conditions compared to the cobblestone morphology observed under perfusion. Scale of embedded image represents 20 µm. Scale of larger lower resolution images represents 200 µm.

## Supplementary Data 11: Analysis of HAEC alignment across perfused IA models

**Figure S11:**
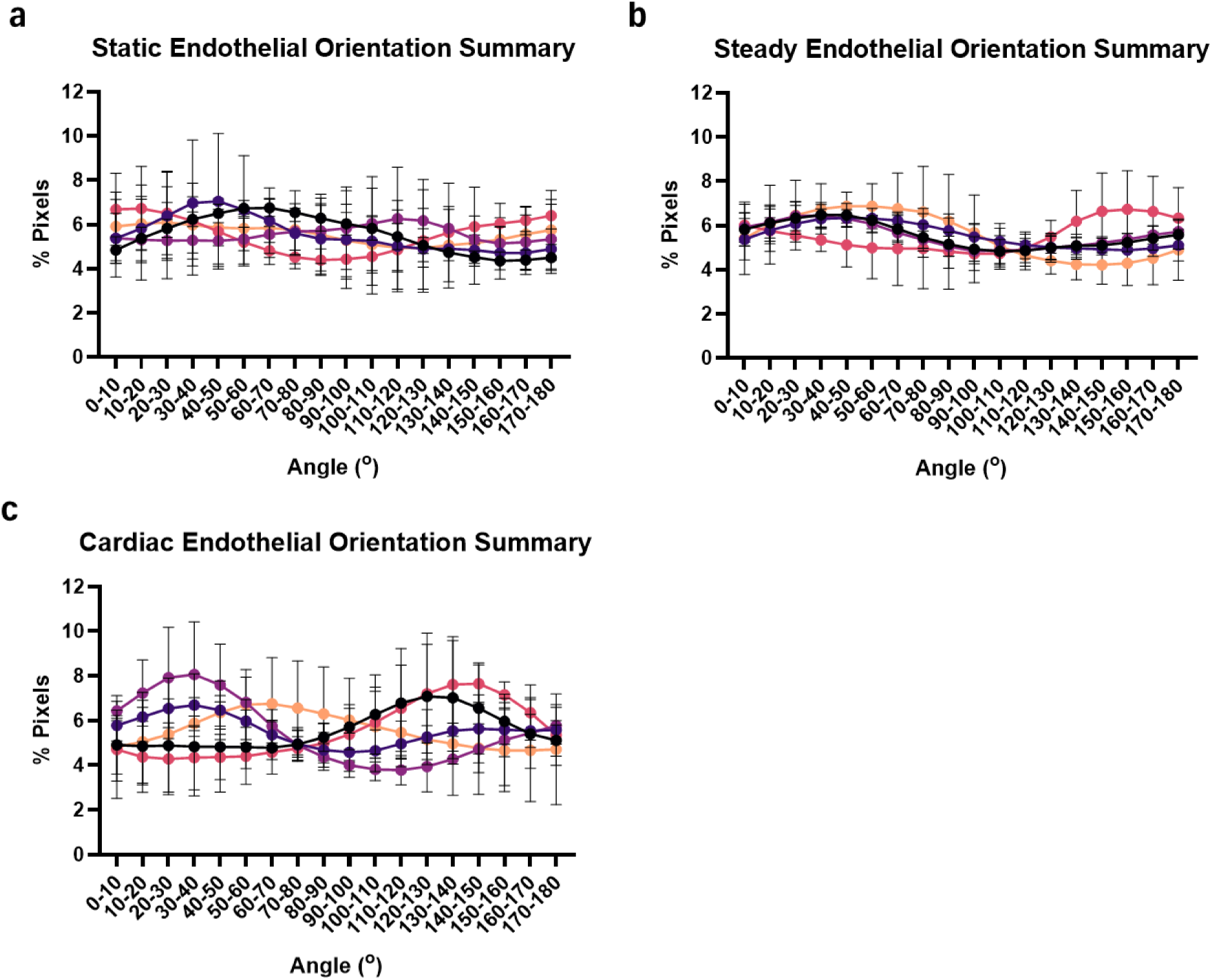
Alignment of human aortic endothelial cells under static, steady and cardiac flow determined using the OrientationJ plugin of ImageJ. (a) CD31 pixel orientation analysis for static cultures. (b) CD31 pixel orientation analysis for aneurysm models exposed to steady flow. (c) CD31 pixel orientation analysis for aneurysm models exposed to cardiac flow waveform.

## Supplementary Data 12: Morphological transition of HAECs under flow

**Figure S12:**
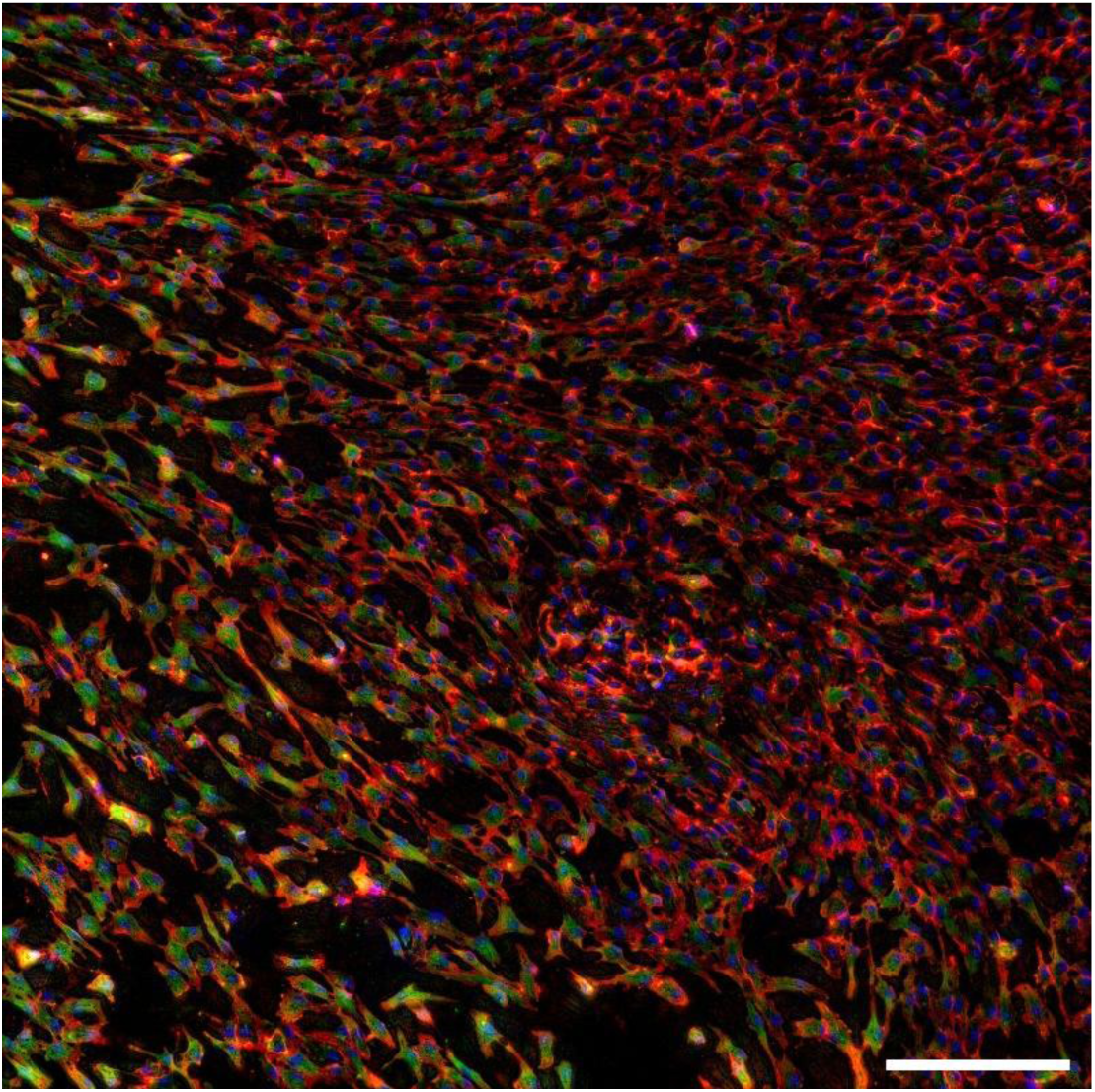
Transition of human aortic endothelial cells (HAECs) from a cobblestone morphology to a sparse and aligned morphology at the aneurysm neck. HAECs stained for F-actin (green), DAPI (blue), CD31 (red) and Hif-1α (magenta). Scale represents 200 µm.

## Supplementary Data 13: Minimal impact of microgrooves on HAEC alignment

**Figure S13:**
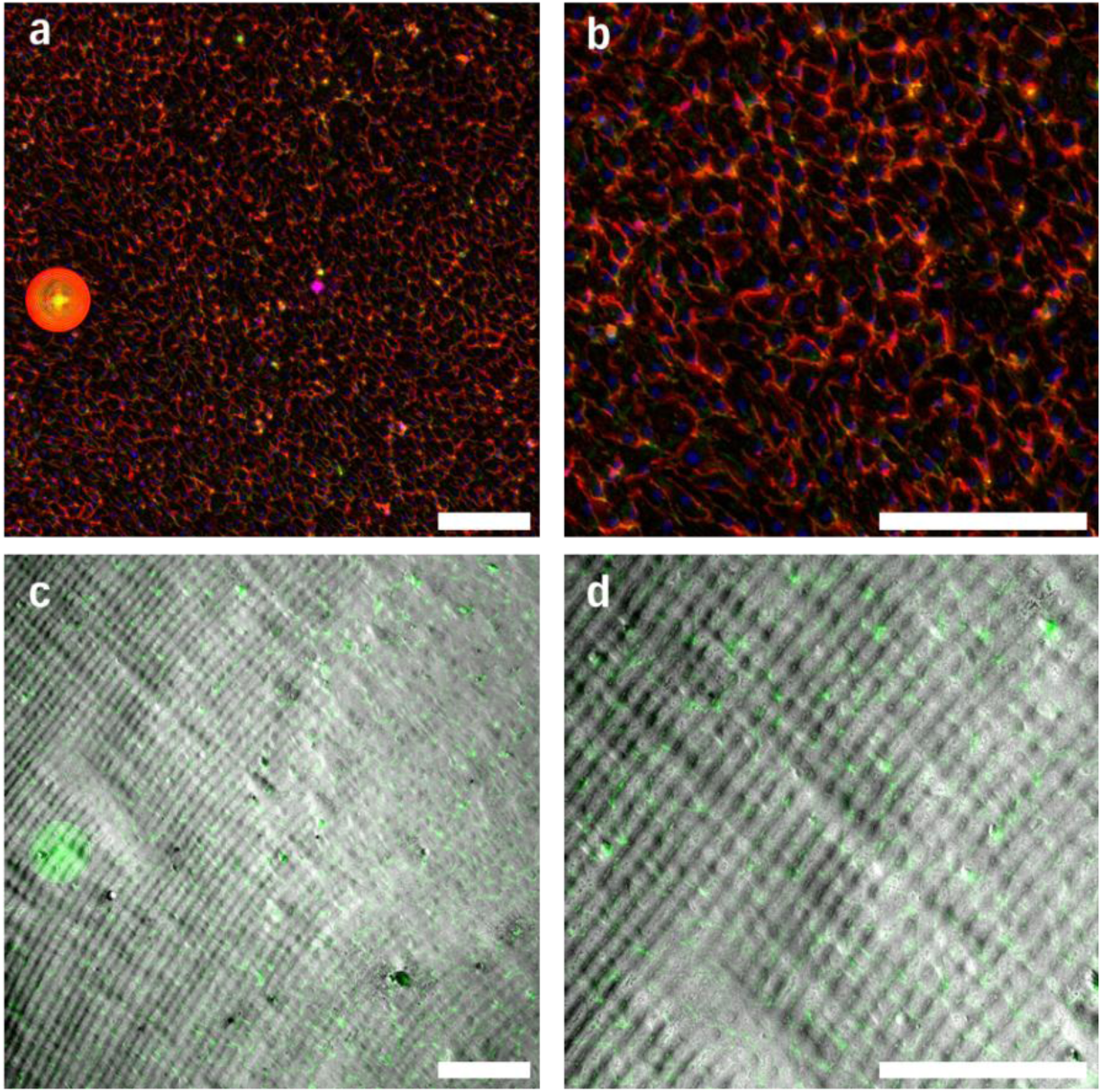
Minimal effect of microgrooves on alignment of human aortic endothelial cells (HAECs) in the left thick-walled lobe of the cardiac flow replicate 2 aneurysm model. (a) Immunofluorescent images of HAECs stained for F-actin (green), DAPI (blue), CD31 (red) and Hif-1α (magenta). (b) Zoomed-in image of (a). (c) Overlay of HAECs stained for F-actin against brightfield image of surface grooves. (d) Zoomed-in image of (c). Scale represents 200 µm.

## Supplementary Data 14: Hif-1α staining with isotypes

**Figure S14:**
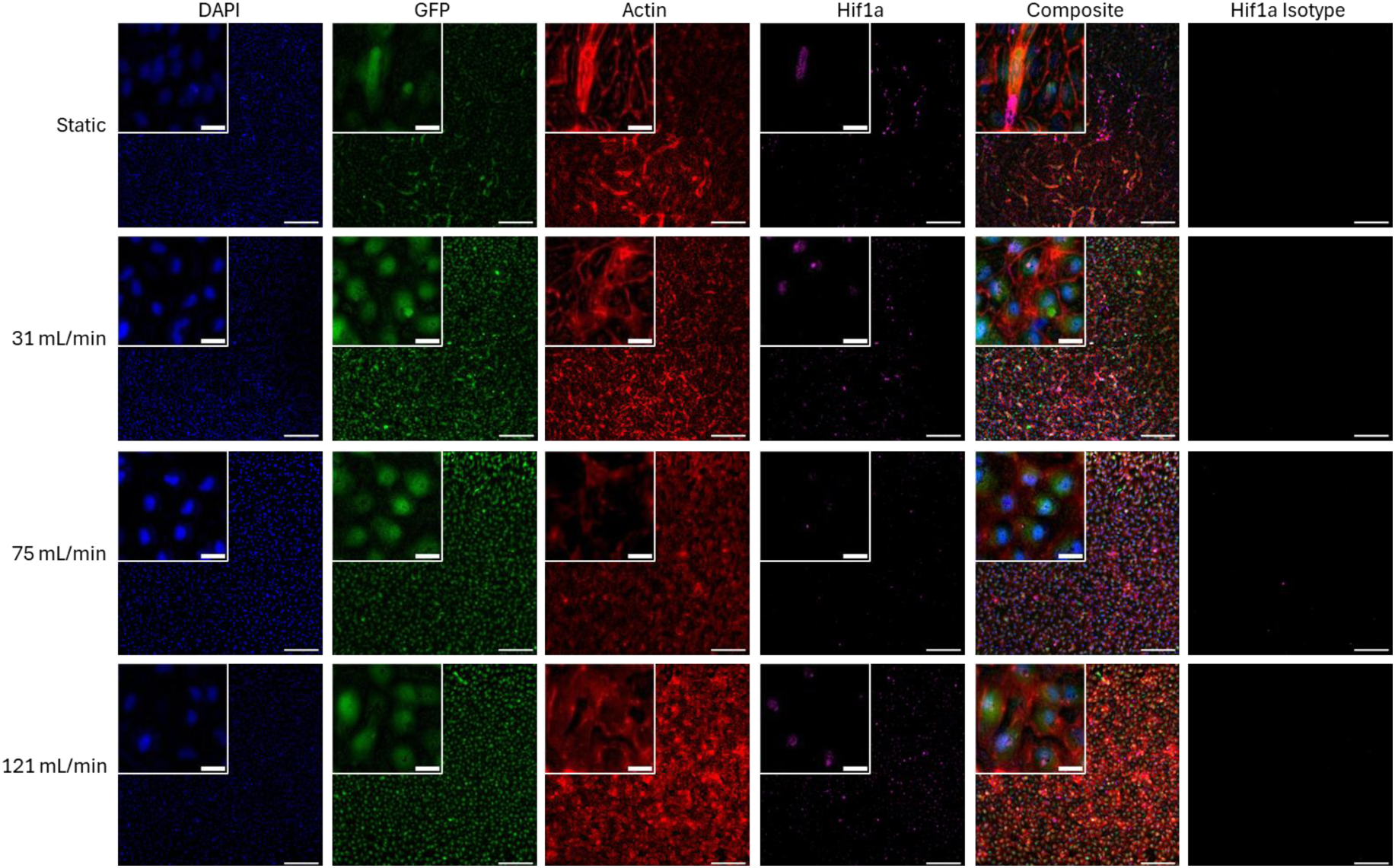
Hif-1α expression and actin staining in human aortic endothelial cells (HAECs) under static and flow conditions. Immunofluorescent images of GFP (green) fluorescing HAECs on PDMS channels stained for DAPI (blue), F-actin (red) and Hif-1α (magenta). Scale of embedded image represents 20 µm. Scale of larger low-resolution image represents 200 µm.

## Supplementary Data 15: Ki67 and eNOS Isotypes

**Figure S15:**
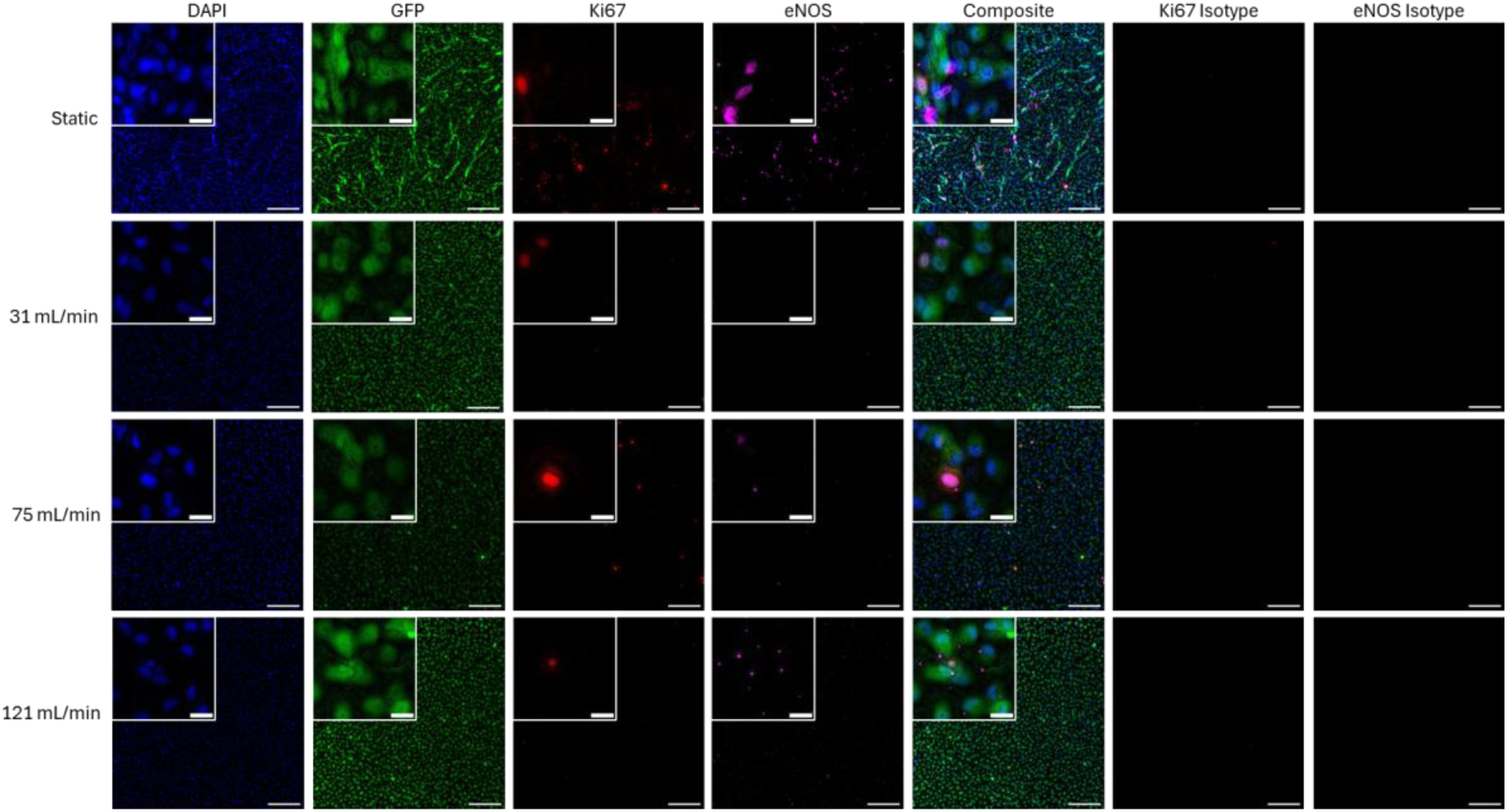
Ki67 and eNOS expression in human aortic endothelial cells (HAECs) under static and flow conditions. Immunofluorescent images of GFP (green) fluorescing HAECs on PDMS channels stained for DAPI (blue), Ki67 (red) and eNOS (magenta). Scale of embedded image represents 20 µm. Scale of larger low resolution image represents 200 µm.

## Supplementary Data 16: Rate of change of EGF and PDGF-AA under perfusion

**Figure S16:**
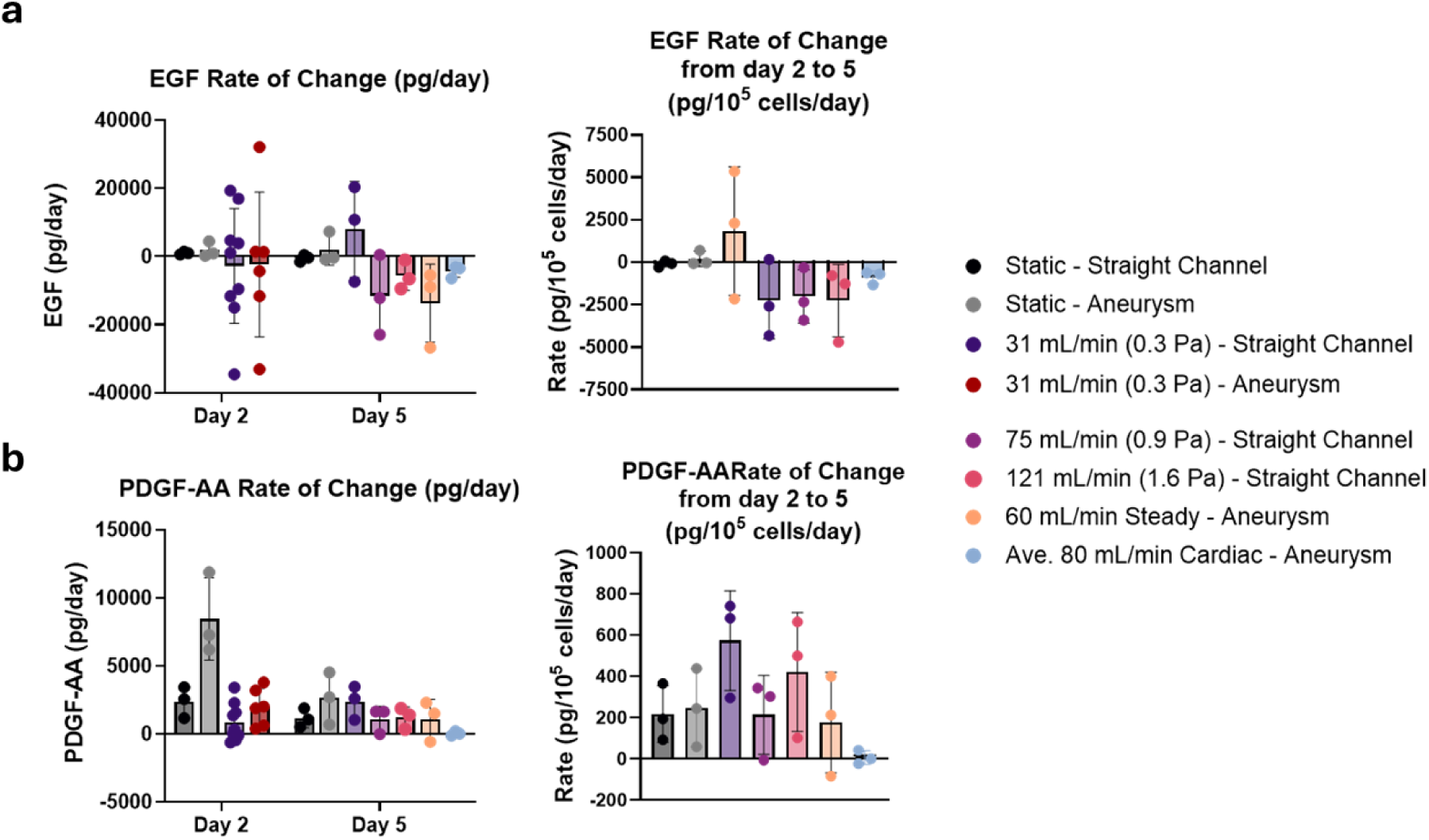
LEGENDplex^TM^ quantification of endothelial growth factors at day 2 and 5. (a-b) volume-corrected and cell density-normalised rate of change of (a) epidermal growth factor (EGF) and (b) Platelet-derived growth factor-AA (PDGF-AA). Cell density normalisation based on quantification of cell density in Figure 3.

## Supplementary Data 17: Original LEGENDplex^TM^ measurements of cytokines and growth factors from perfused channels

Original measurements from LEGENDplex^TM^ analysis before volume-correction. Run-specific LOQ’s have been marked with a dotted line with corresponding analyte measurements represented by either a closed (●) or open (○) circle, or a triangle (▲). LOQs that were far below the observed measurements were omitted for clarity.

FGF-b degraded quickly with incubation time, with an estimated half-life of 0.43 days or 10.4 hours assuming first-order decay between day zero and day two. This is consistent with existing literature on FBF-b stability ^102,103^. Limited FGF-b was therefore available for cellular consumption by day five across all conditions.

Measured VEGF concentration in media controls were relatively consistent and did not show clear degradation with incubation duration as was expected ^104–106^. This variability could reflect preparation differences, decay with storage duration, instability with freezing, VEGF stabilisation by serum proteins, or assay detectability differences ^107,108^. The slight increase in VEGF concentration measured at day five in media controls could reflect the addition of fresh media at day two.

**Figure S17:**
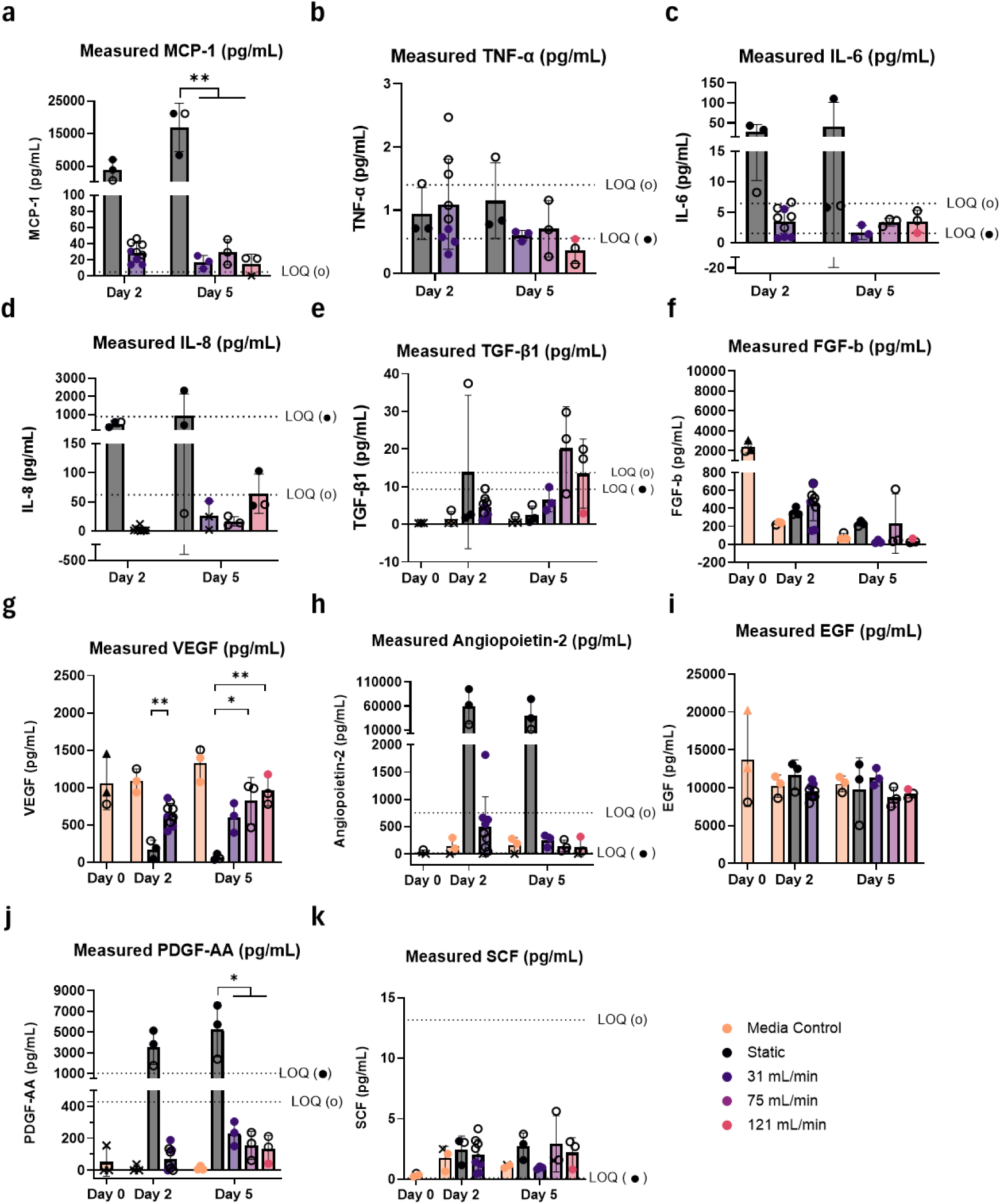
LEGENDplex^TM^ quantification of endothelial cytokines and growth factors at day 2 and 5 in straight channel perfusion experiments. Raw measurements of (a) Monocyte chemoattractant protein-1 (MCP-1), (b) Tumour Necrosis Factor-alpha (TNF-α), (c) Interleukin-6 (IL-6), (d) Interleukin-8 (IL-8), (e) Transforming growth factor-β1 (TGF-β1), (f) Fibroblast growth factor-basic (FGF-b), (g) Vascular endothelial growth factor (VEGF), (h) Angiopoietin-2, (i) epidermal growth factor (EGF), (j) Platelet-derived growth factor-AA (PDGF-AA) and (k) Stem-cell factor (SCF).

## Supplementary Data 18: Original LEGENDplex^TM^ measurements of cytokines and growth factors from perfused IA models

**Figure S18:**
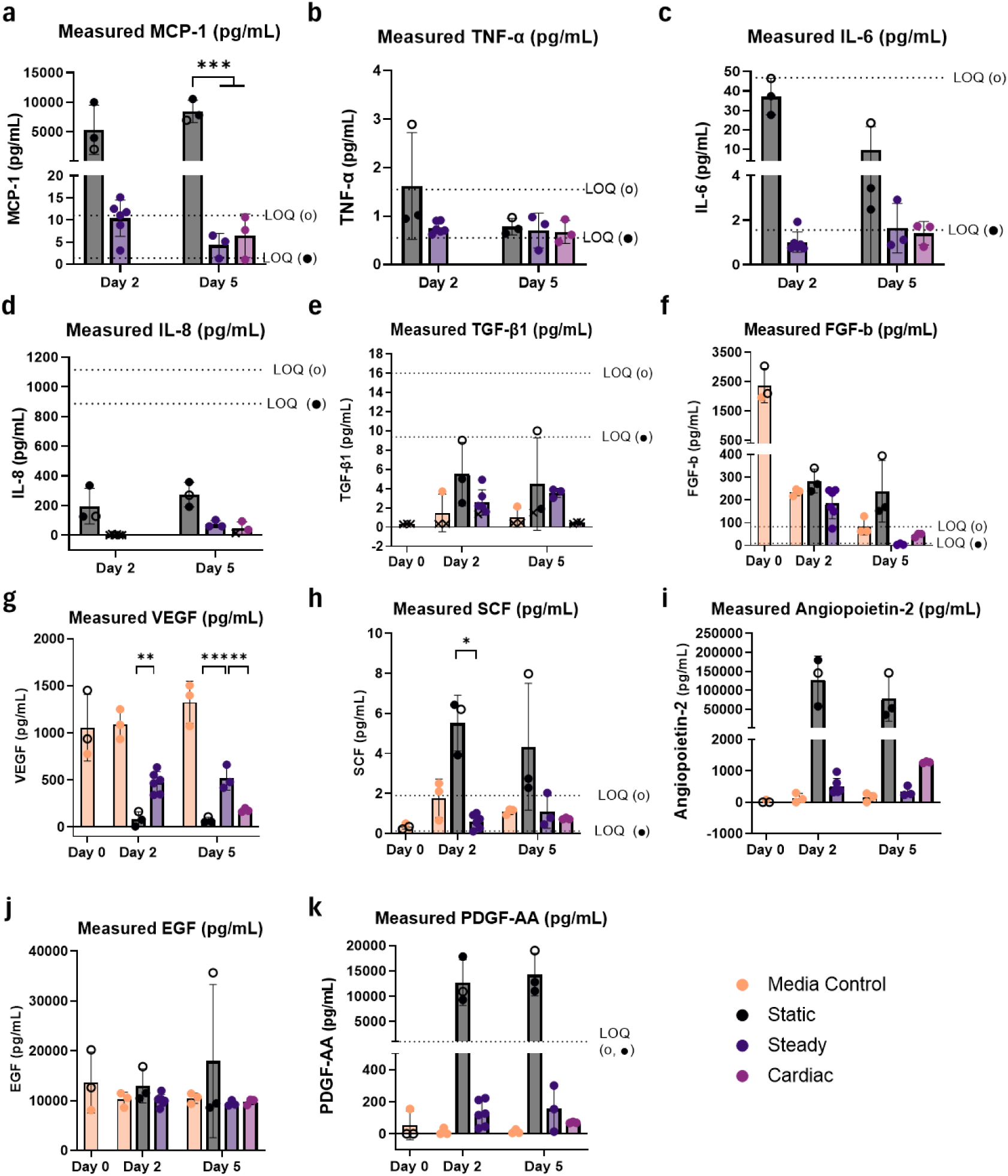
LEGENDplex^TM^ quantification of endothelial cytokines and growth factors at day 2 and 5 in patient-specific IA model perfusion experiments. Raw measurements of (a) Monocyte chemoattractant protein-1 (MCP-1), (b) Tumour Necrosis Factor-alpha (TNF-α), (c) Interleukin-6 (IL-6), (d) Interleukin-8 (IL-8), (e) Transforming growth factor-β1 (TGF-β1), (f) Fibroblast growth factor-basic (FGF-b), (g) Vascular endothelial growth factor (VEGF), (h) Angiopoietin-2, (i) epidermal growth factor (EGF), (j) Platelet-derived growth factor-AA (PDGF-AA) and (k) Stem-cell factor (SCF).

## Supplementary Data 19: Example of MCP-1 cytokine calculations

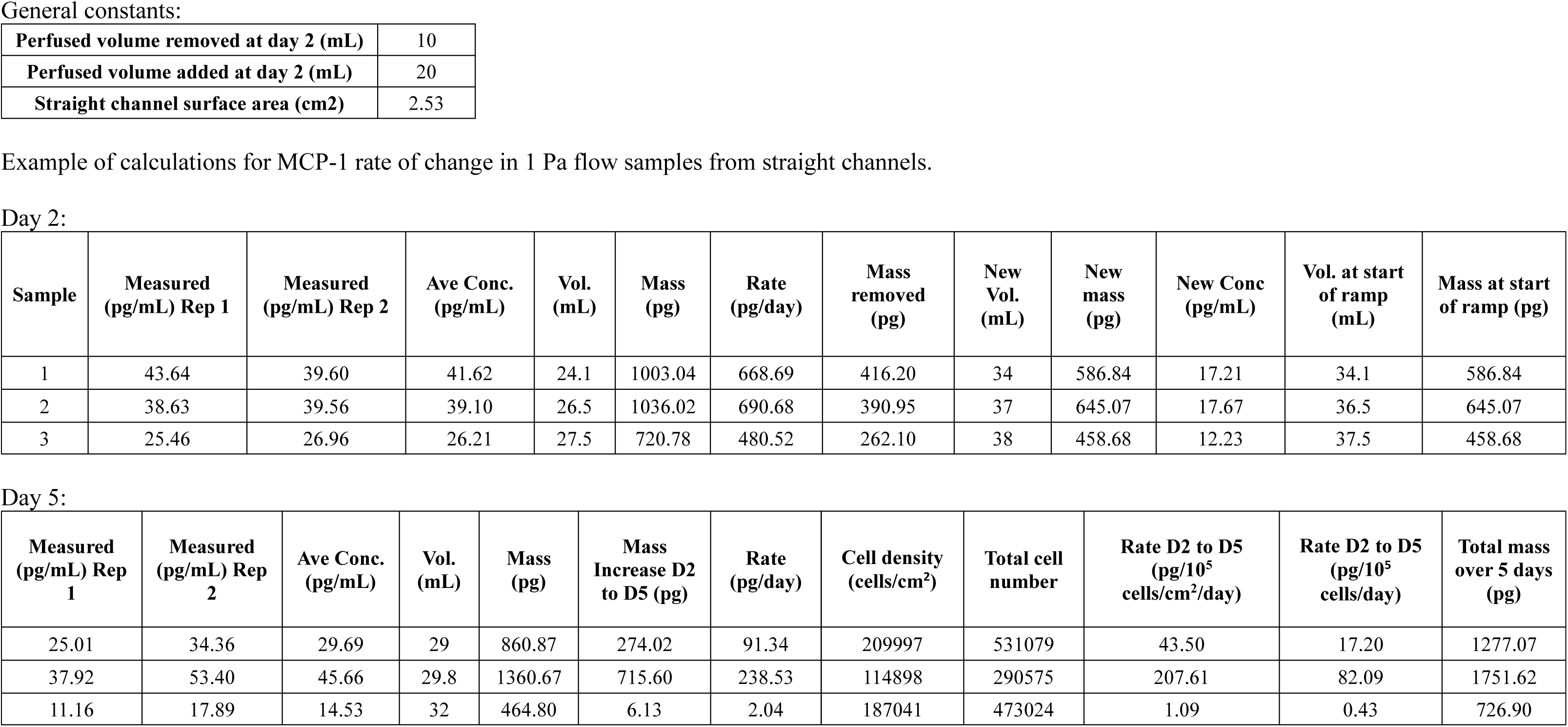

## Supplementary Data 20: Example of FGF-b growth factor calculations

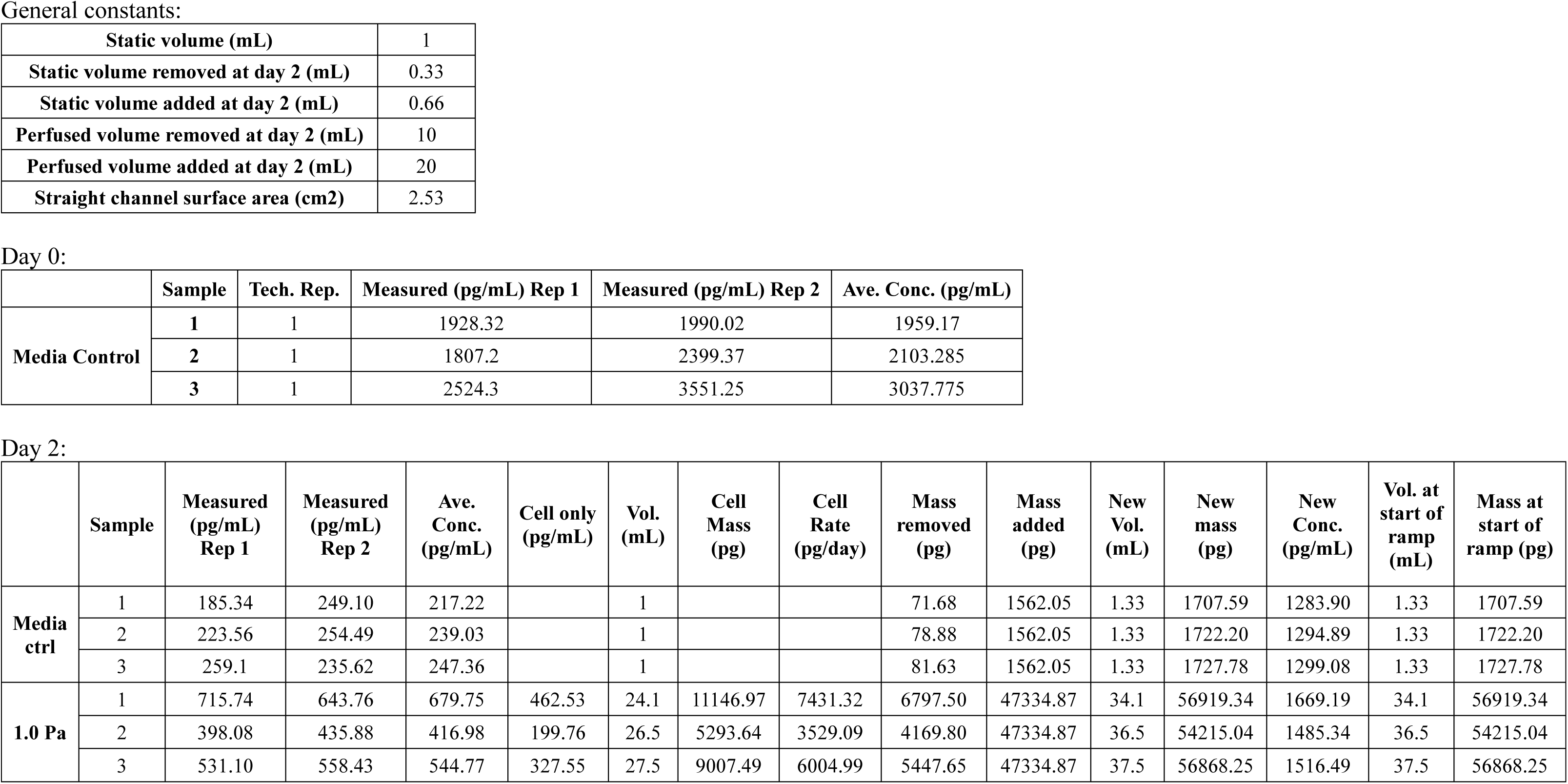
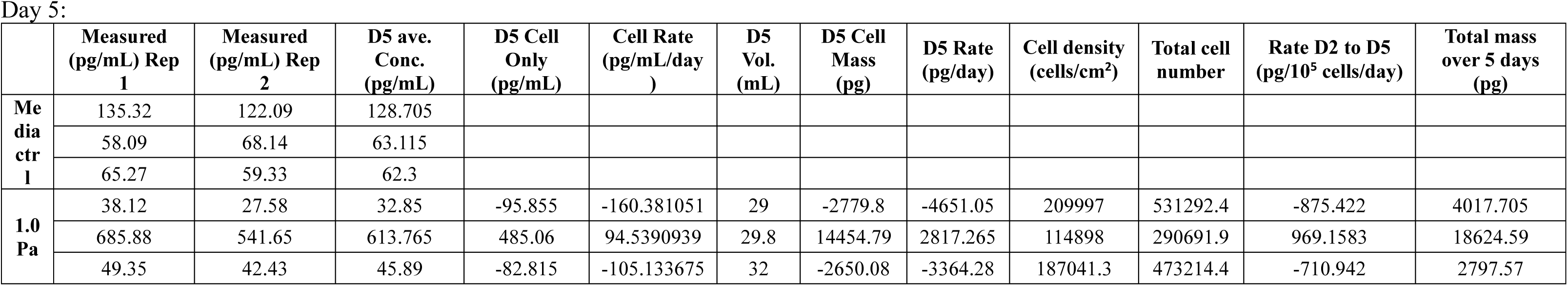

